## Supporting Tables for "The N-terminus of varicella-zoster virus glycoprotein B has a functional role in fusion"

**S1 Table.** Cryo-EM data collection parameters for the native full-length VZV gB (EMDB 22629), and the gB-93k (EMDB 22519) and gB-SG2 (EMDB 22520) complexes.

| Parameter | Structure | | |
|---|---|---|---|
| | **Native VZV gB** | **gB-93k Fab** | **gB-SG2 Fab** |
| **Data collection and processing** | | | |
| Microscope | Titan Krios (FEI) | Tecnai F20 (FEI) | Tecnai F20 (FEI) |
| Magnification (1000X) | 130 | 29 | 29 |
| Voltage (kV) | 300 | 200 | 200 |
| Energy filter slit width (eV) | 20 | N/A | N/A |
| Detector | Gatan K2 Summit | Gatan K2 Summit | Gatan K2 Summit |
| Defocus range (µm) | 1.5-2.0 | 1.8-3 | 1.8-3 |
| Pixel size (Å) | 1.06 | 1.283 | 1.283 |
| Exposure time (s) | 12 | 15 | 15 |
| Frames | 60 | 75 | 75 |
| Electron dose ($e^-$/s) | 7.5 | 7.8 | 7.8 |
| Electron exposure rate ($e^-$/Å$^2$/s) | 1.335 | 4.2 | 4.2 |
| Total electron exposure ($e^-$/Å$^2$) | 16.02 | 63 | 63 |
| Symmetry imposed | 3 | 3 | 3 |
| Micrographs collected (no.) | 10,241 | 649 | 795 |
| Final particle images (no.) | 349,207 | 22,976 | 25,670 |
| Map resolution (Å) | 3.9 | 7.3 | 9.0 |
| FSC threshold | 0.143 | 0.143 | 0.143 |
| **Refinement** | | | |
| Initial model used (PDB code) | 6VLK[A] | N/A | N/A |
| Model resolution (Å) | 3.9 | N/A | N/A |
| FSC threshold | 0.143 | N/A | N/A |
| Model composition | | N/A | N/A |
| Chains | 6 | N/A | N/A |
| Non-hydrogen atoms | 14,409 | N/A | N/A |
| Protein residues | 1,752 | N/A | N/A |
| **Validation** | | | |
| B factors (Å$^2$) | | N/A | N/A |
| Protein (min/max/mean) | 42/164/72 | N/A | N/A |
| Protein (min/max/mean) | 59/151/93 | N/A | N/A |
| R.M.S. deviations | | N/A | N/A |
| Bond lengths (Å) | 0.006 | N/A | N/A |
| Bond angles (°) | 0.851 | N/A | N/A |
| Validation | | N/A | N/A |
| MolProbity score | 1.7 | N/A | N/A |
| Clashscore | 15 | N/A | N/A |
| Poor rotamers (%) | 0.4 | N/A | N/A |
| Ramachandran plot | | N/A | N/A |
| Favored (%) | 98 | N/A | N/A |
| Allowed (%) | 2 | N/A | N/A |
| Disallowed (%) | 0 | N/A | N/A |

[A] X-ray crystallography data for VZV gB.

**S2 Table.** X-ray data collection and structure refinement for VZV gB (PDB 6VLK).

| **Parameter** | |
|---|---|
| **X-ray Data** | |
| Beamline | APS 17-ID |
| Space group | R32 |
| Cell dimensions: a, b, c | 118.318, 118.318, 749.026 |
| (Å), α, β, γ (°) | 90, 90, 120 |
| Resolution (Å) | 101.52-2.45 (2.59-2.45)[A] |
| $R_{merge}$ | 0.073 (0.516)[A] |
| I/σ | 19.8(3.7)[A] |
| Completeness (%) | 100.0(100.0)[A] |
| Redundancy | 9.8(10.0)[A] |
| **Refinement** | |
| Model Resolution(Å) | 50.2-2.4 |
| Number of reflections | 74911 |
| $R_{work}/R_{free}$[B] | 0.1845/0.23 |
| Model Composition | |
| Protein | 9381 |
| Glycan | 214 |
| Water | 171 |
| **Validation** | |
| B factors (Å$^2$) | |
| Protein (min/max/mean) | 7/155/44 |
| Glycan (min/max/mean) | 24/145/43 |
| R.M.S. deviations | |
| Bond lengths (Å) | 0.01 |
| Bond angles (°) | 1.14 |
| Validation | |
| MolProbity score | 1.69 |
| Clashscore | 3 |
| Poor rotamers (%) | 4.6 |
| Ramachandran plot | |
| Favored (%) | 98 |
| Allowed (%) | 2 |
| Disallowed (%) | 0 |

[A]Outer-shell values are given in parentheses.

[B]The *Rfree* test set was composed of 5% randomly chosen reflections.

**S3 Table.** Amino acid residues and color code for each domain in VZV gB.

| Residues | Domain | Color |
|---|---|---|
| 115-136 | IV | Orange |
| 137-147 | Linker: II-IV | Hot Pink |
| 148-159 | II | Green |
| 160-368 | I | Cyan |
| 369-464 | II | Green |
| 465-502 | II? | |
| 503-510 | Linker: II-III | Hot Pink |
| 511-569 | III | Yellow |
| 570-681 | IV | Orange |
| 682-736 | V | Red |

**S4 Table.** N-linked glycosylation sites identified in VZV and herpesvirus gB orthologues.

| Domain | VZV | | HSV | PRV | HCMV | EBV |
|---|---|---|---|---|---|---|
| | Predicted | Observed | | | | |
| DII/IV linker | N147 | N147[B/C] | N141 | N154 | X | N76 |
| DI | N257 | N257[A/B/C] | X – N255 | N264 | X | X |
| DII | N435 | N435[B/C] | N430 | N444 | N409 | X |
| DII no structure | N479 | | | | | |
| DII/III linker | N503 | N503[C] | | | | |
| DIII | N557 | | | | | |
| DIV | N620 | N620[A/B] | | N636 | N586 | N563 |
| DV | N686 | N686[A/B/C] | | | | |

[A] Identified in the gB ectodomain X-ray crystal structure
[B] Density observed in Cryo-EM map
[C] Identified by Orbitrap Mass Spectrometry

**S5 Table.** Conserved cysteine bonds in herpesvirus gB orthologues.

| Domain Location | Cysteine Residue Locations in gB | | | | |
|---|---|---|---|---|---|
| | **VZV** | **HSV** | **PRV** | **HCMV** | **EBV** |
| DIV | 122-584 | 116-573 | 129-603 | 94-551 | 51-528 |
| DII/IV linker to DIII | 139-540 | 133-529 | 146-559 | 111-507 | 68-484 |
| DI | 213-277 | 207-271 | 220-284 | 185-250 | 141-206 |
| DII | 369-417 | 364-412 | 377-426 | 344-391 | 295-342 |
| DIV | 608-645 | 596-633 | 625-661 | 574-611 | 551-588 |

**S6 Table.** Amino acid residues from X-ray crystallography data for the herpesvirus gB orthologues used to calculate amino acid identities and RMSD.

| **Domain** | **Amino acid residues within each domain** | | | |
|---|---|---|---|---|
| | **HSV (2GUM[A])** | **PRV (6ESC[A])** | **HCMV (5CXF[A])** | **EBV (3FVC[A])** |
| Complete Structure | 111-724 | 122-750 | 87-695 | 44-679 |
| I | 154-363 | 167-376 | 133-343 | 89-294 |
| II | 142-153/364-458 | 155-166/377-473 | 121-132/344-438 | 77-88/295-389 |
| III | 492-558 | 522-589 | 476-536 | 448-513 |
| IV | 111-130/559-669 | 122-143/590-695 | 87-108/537-647 | 44-65/514-624 |
| V | 670-724 | 696-750 | 648-695 | 625-679 |

[A] Protein Data Bank accession number

**S7 Table.** Amino acid identities of VZV gB derived from structure-based alignments with herpesvirus gB orthologues.

| **Domain** | **Amino acid identity (similarity) [%] compared to VZV** | | | |
|---|---|---|---|---|
| | **HSV** | **PRV** | **HCMV** | **EBV** |
| Complete Structure | 50 (71) | 61 (77) | 30 (51) | 29 (50) |
| I | 48 (64) | 61 (75) | 28 (44) | 27 (43) |
| II | 46 (74) | 50 (70) | 32 (53) | 22 (46) |
| III | 54 (73) | 57 (75) | 33 (61) | 26 (55) |
| IV | 50 (70) | 58 (73) | 29 (50) | 40 (56) |
| V | 63 (89) | 78 (92) | 32 (57) | 27 (55) |

**S8 Table.** RMSD of VZV gB derived from structure-based alignments with herpesvirus gB orthologues.

| Domain | RMSD (Å)$^{A}$ compared to VZV gB | | | | | | | |
|---|---|---|---|---|---|---|---|---|
| | HSV | | PRV | | HCMV | | EBV | |
| Complete Structure | 0.987 (473) | 1.643 (574) | 0.855 (337) | 2.880 (578) | 1.268 (136) | 7.154 (558) | 1.154 (226) | 10.025 (546) |
| I | 0.635 (195) | 1.391 (203) | 0.593 (198) | 1.008 (209) | 1.054 (108) | 3.657 (205) | 1.084 (156) | 2.536 (204) |
| II | 0.635 (95) | 15.115 (107) | 0.779 (91) | 15.141 (107) | 0.863 (80) | 13.179 (101) | 0.827 (70) | 11.753 (99) |
| III | 0.548 (67) | 0.548 (67) | 0.709 (60) | 1.687 (68) | 0.847 (61) | 0.847 (61) | 0.936 (60) | 1.641 (66) |
| IV | 1.022 (96) | 6.133 (125) | 0.892 (94) | 6.904 (123) | 0.978 (80) | 4.724 (116) | 1.017 (80) | 8.040 (110) |
| V | 0.649 (55) | 0.649 (55) | 0.531 (55) | 0.531 (55) | 1.149 (29) | 4.260 (48) | 1.157 (38) | 3.032 (51) |

[A] Root mean squared deviation (RMSD) calculated using Matchmaker (Chimera). The two values correspond to the trimmed amino acids versus all of the amino acids in the alignment used by Matchmaker.

**S9 Table.** Key reagents and resources.

| REAGENT or RESOURCE | SOURCE | IDENTIFIER |
|---|---|---|
| **Antibodies** | | |
| Human mAb 93k | This paper | |
| Mouse mAb SG2-2E6 | GeneTex | GTX38718 |
| Rabbit polyclonal antibody 746-868 | [7] | |
| Mouse mAb IE62 | EMD Millipore | MAB8616 |
| Mouse mAb gE | EMD Millipore | MAB8612 |
| VZV mouse mixed mAb | Meridian Life Sciences | C05108MA |
| Mouse anti-V5 tag | Bio-Rad | MCA1360 |
| Biotinylated goat anti-mouse IgG (H+L) | Vector Labs. Inc | BA-9200 |
| Donkey anti-mouse Alexa Fluor 555 | Life Technologies | A31570 |
| Donkey anti-human Alexa Fluor 488 | Life Technologies | A11013 |
| ECL Sheep anti-mouse IgG, Horseradish peroxidase linked whole antibody | GE Healthcare UK Ltd | NA931V |
| ECL Donkey anti-rabbit IgG, Horseradish peroxidase linked whole antibody | GE Healthcare UK Ltd | NA934V |
| ECL Sheep anti-human IgG, Horseradish peroxidase linked whole antibody | GE Healthcare UK Ltd | NA933V |
| **Bacterial and Virus Strains** | | |
| pOka BAC derived (pPOKA-DX) | [8] | |
| GS1783 | [9] | |
| pOka-TK-GFP | [10] | |
| pOka-TK-GFP gB-STEVV5 | This paper | |
| pOka-TK-GFP gB-TEVV5 | This paper | |
| pOka-TK-GFP gB-TEVV5 gB[S110A] | This paper | |
| pOka-TK-GFP gB-TEVV5 gB[Q111A] | This paper | |
| pOka-TK-GFP gB-TEVV5 gB[D112A] | This paper | |
| pOka-TK-GFP gB-TEVV5 gB[$^{109}$AAAA$^{112}$] | This paper | |
| **Chemicals, Peptides, and Recombinant Proteins** | | |
| Anti-V5 Agarose Affinity Gel | Sigma | A7345-1ML |
| Bovine Serum Albumin (IgG Free, Protease Free) | Jackson ImmunoResearch | 001-000-162 |
| Minimal essential medium | Corning cellgro | 10-010-CV |
| F-12K nutrient mixture Kaighn's modification | Invitrogen | 21127-022 |
| Optimem +Glutamax | Gibco | 51985-034 |
| Fetal bovine serum | Gibco | 26140-079 |
| Penicillin /Streptomycin | Gibco | 15140-122 |
| Amphotericin B | Corning cellgro | 30-003-CF |
| Nonessential amino acids | Corning cellgro | 25-025-CI |
| Puromycin | Invitrogen | A11138-03 |

| | | |
|---|---|---|
| BstZ171 | New England BioLabs Inc. | R3594 |
| NaeI | New England BioLabs Inc. | R0190L |
| HindIII | New England BioLabs Inc. | R0104L |
| AgeI | New England BioLabs Inc. | R0552L |
| SpeI | New England BioLabs Inc. | R0133L |
| NotI | New England BioLabs Inc. | R0189L |
| KpnI | New England BioLabs Inc. | R0142L |
| XmaI | New England BioLabs Inc. | R0180L |
| NdeI | New England BioLabs Inc. | R0111L |
| AccuPrime *Pfx* DNA Polymerase | Invitrogen | 12344024 |
| KOD Extreme Hot Start DNA Polymerase | EMD Millipore | 71975-3 |
| Lipofectamine 2000 | Invitrogen | 11668-019 |
| Ampicillin | Sigma | A9518-100G |
| Kanamycin | Sigma | K4378 |
| LB (Miller's) Agar | Growcells | MBPE-3060 |
| LB (Miller's) Broth | Growcells | MBPE-1050 |
| PBS (Phosphate buffered saline) | Corning cellgro | 21-040-CV |
| DPBS (Dulbecco's phosphate buffered saline) | Corning cellgro | 21-030-CV |
| Tris Base | Fisher Scientific | BP152-5 |
| Sodium chloride | Fisher Scientific | S271-10 |
| Potassium chloride | Fisher Scientific | BP366-500 |
| Magnesium chloride | Fisher Scientific | M33-500 |
| Sodium deoxycholate | ICN Biomedical Inc. | 804312 |
| IGEPAL CA-630 | Sigma | I3021-100ML |
| Triton X-100 | Sigma | T-9284 |
| Tobacco etch virus protease | In house | |
| Tris buffered saline (TBS; pH7.4) | Scy Tek | TBS500 |
| Amphipol 8-35 | Anatrace | A835 |
| Lauroylsarcosine | Sigma | L9150 |
| Bio-Beads SM-2 | Bio-Rad | 152-3920 |
| Phosphotungstic acid | Ted Pella | 19402 |
| EDTA | Sigma | E9884 |
| Sucrose | Sigma | S7903-1KG |
| L(+)-glutamic acid monosodium salt monohydrate | Sigma | G1626-100G |
| Paraformaldehyde (4%) in PBS | Boston Bioproducts | K06J101 |
| BD Cytofix/Cytoperm Plus | BD Biosciences | 555028 |

| | | |
|---|---|---|
| Hoechst 33342 | ThermoFisher Scientific | H-3570 |
| Native PAGE Sample Buffer | Novex | BN20032 |
| Native PAGE running buffer 20X | Novex | BN2001 |
| Native PAGE 20X cathode buffer additive | Novex | BN2002 |
| Native PAGE 3-12% Bis Tris Gel | Novex | BN2011BX10 |
| Laemmli Sample Buffer 2X | Bio-Rad | 161-0737 |
| 2-mercaptoethanol | Sigma | M7522-100ML |
| Mini Protean TGX Gels 4-20% | Bio-Rad | 456-1094 |
| NativeMark Protein Std. | Invitrogen | 57030 |
| Novex Sharp Pre-Stained Protein Standards | Invitrogen | 57318 |
| Dimethyl pimelimidate dihydrochloride | Sigma | D8388 |
| Protein A Plus UltraLink Resin | Thermo Scientific | 53142 |
| Fluoromount-G | SouthernBiotech | 0100-01 |
| Boric acid | Sigma | B-0252 |
| Ethanolamine | Sigma | 398136-500ML |
| Ethane | Airgas | ET R80 |
| L-(+)-Arabinose | Sigma | A3256-100G |
| Agarose LE | AccuFlow | EK2808 |
| Membrane permeable coelenterazine-H | Nanolight Technology | 3012-10 |
| Fast Red TR Salt hemi (zinc chloride) | Sigma | 368881-25G |
| Naphthol AS-MX phosphate | Sigma | N4875-500MG |
| Alkaline phosphatase-conjugated Streptavidin | Jackson ImmunoResearch | 016-050-084 |
| **Experimental Models: Cell Lines** | | |
| MeWo | ATCC | HTB-65 |
| CHO DSP1 | [11] | |
| Mel-DSP2 | [11] | |
| **Oligonucleotides (All sequences are 5' to 3')** | | |
| **S/TEVV5** | | |
| gB-AgeI<br>CTTTTTTGCGTACCGGTACGTGC | [12] | Elim Bio |
| gB931<br>[PHOS]CACCCCCGTTACATTCTCGGTGCG | [12] | Elim Bio |
| gB-V5<br>[PHOS]GGTAAGCCTATCCCTAACCCTCTCCTCGGTCTCGATTCTACGTAAATAGCCAGGGGGTTT | [12] | Elim Bio |
| M13R<br>CACCCCCGTTACATTCTCGGTGCG | Invitrogen | |
| gB-Cterm-S-tag<br>[PHOS]GCTGTCCATGTGCTGGCGTTCGAATTTAGCAGCAGCGGTTTCTTTCACCCCCGTTACATTCTCGG | [12] | Elim Bio |

| | | |
|---|---|---|
| gB-link_TEV_link<br>[PHOS]*GGCGGCGGGGGCGGG*<u>GAGAATCTTTATTTTCAGG</u>*GCGGGGGCGG*GGGGTAAGCCTATCCCTAACCC | [12] | Elim Bio |
| ΔS-tag-sense<br>[PHOS]GGCGGCGGGGGCGGGGAGATTC | [12] | Elim Bio |
| ΔS-tag-antisense<br>[PHOS]CACCCCCGTTACATTCTCGGTG | [12] | Elim Bio |
| [31]F56625-56645<br>AGGTATAGGCAGTTCCCACGG | [13] | Elim Bio |
| [31]R59697-59717<br>TTTCATTGAGACTTGAAGCGC | [13] | Elim Bio |
| **gB KSQD 109/110/111/112** | | |
| pCAGGs-gB-NotI-sense<br>AAAGAATTCGCGGCCGCTGACCG | This paper | Elim Bio |
| gB-93k-sense<br>[PHOS]CAGGACGCCGAAACAAAACCCACGTTTTACG | This paper | Elim Bio |
| K109R-sense<br>[PHOS]CGGTCCCAGGACGCCGAAACAAAACC | | |
| Q111A-sense<br>[PHOS]GCGGACGCCGAAACAAAACCCACGTTTTACG | This paper | Elim Bio |
| K112A-sense<br>[PHOS]CAGGCTGCCGAAACAAAACCCACGTTTTACG | This paper | Elim Bio |
| Q111A-K112A sense<br>[PHOS]GCGGCTGCCGAAACAAAACCCACGTTTTACG | This paper | Elim Bio |
| pCAGGs-gB-KpnI-antisense<br>TCTCTGAAACGGGAATTGGTACC | This paper | Elim Bio |
| gB-93k-antisense<br>GGACTTGTGTATAGCTTCTCTGATTTCATCACC | This paper | Elim Bio |
| K109A-antisense<br>[PHOS]GGACGCGTGTATAGCTTCTCTGATTTCATCACC | This paper | Elim Bio |
| H108-antisense<br>[PHOS]GTGTATAGCTTCTCTGATTTCATCAC | This paper | Elim Bio |
| S110A-antisense<br>[PHOS]AGCCTTGTGTATAGCTTCTCTGATTTCATCACC | This paper | Elim Bio |
| K109A-S110A-antisense<br>[PHOS]AGCCGCGTGTATAGCTTCTCTGATTTCATCACC | This paper | Elim Bio |
| | This paper | Elim Bio |
| **Recombinant DNA** | | |
| pCAGGs-VZVgB | [14] | |
| pME18s | [14] | |
| pME18s-gH[TL] | [14] | |
| pME18s-gH[V5] | [10] | |
| pCDNA3.1(+) | Invitrogen | |
| pCDNA3.1-gL | [15] | |
| pBud-gE/gI | [16] | |

| | | |
|---|---|---|
| pCAGGs-gB[K109A] | This paper | |
| pCAGGs-gB[K109R] | This paper | |
| pCAGGs-gB[S110A] | This paper | |
| pCAGGs-gB[Q111A] | This paper | |
| pCAGGs-gB[D112A] | This paper | |
| pPOKA-TK-GFP gB-STEVV5 | This paper | |
| pPOKA-TK-GFP gB-TEVV5 | [12] | |
| pPOKA-TK-GFP gB-TEVV5 gB[K109A] | This paper | |
| pPOKA-TK-GFP gB-TEVV5 gB[K109R] | This paper | |
| pPOKA-TK-GFP gB-TEVV5 gB[S110A] | This paper | |
| pPOKA-TK-GFP gB-TEVV5 gB[Q111A] | This paper | |
| pPOKA-TK-GFP gB-TEVV5 gB[D112A] | This paper | |
| **Software and Algorithms** | | |
| SerialEM | [17] | http://bio3d.colorado.edu/SerialEM/ |
| Relion v3.0 | [18, 19] | https://bitbucket.org/scheres/relion-3.0_beta/src/master/ |
| ResMap v1.95 | [20] | https://sourceforge.net/projects/resmap-latest/ |
| UCSF Chimera v1.13.1 | [21] | http://www.cgl.ucsf.edu/chimera/ |
| ModelZ | [22] | https://cryoem.slac.stanford.edu/ncmi/resources/software/modelz |
| Phenix | [23] | https://www.phenix-online.org/ |
| WinCoot | [24] | http://bernhardcl.github.io/coot/ |
| RaptorX | [25] | http://raptorx.uchicago.edu/ |
| SWISS-MODEL | [26] | https://swissmodel.expasy.org/ |
| CHARMM-GUI | [27] | http://www.charmm-gui.org/ |
| Segger v1.9.5 | [28] | https://cryoem.slac.stanford.edu/ncmi/resources/software/segger |
| CellQuest Pro | BD | |

| | | |
|---|---|---|
| FlowJo | TreeStar | |
| Prism 8 | GraphPad Software, Inc. | |
| Staden | [29] | http://staden.sourceforge.net |
| GeneDoc v2.7 | Nicholas and Nicholas, 1997 | https://genedoc.software.informer.com/download/ |
| FiJi (ImageJ 1.52i) | NIH, USA | http://imagej.nih.gov/ij |
| Illustrator CS6 | Adobe | |
| Photoshop CS6 | Adobe | |
| AxioVision | Zeiss | |
| **Other** | | |
| NativePAGE 2-12% Bis-Tris Gel | Invitrogen | BN2011BX10 |
| Mini-PROTEAN TGX Gels | Bio-Rad | 456-1094 |
| Superose-6 Increase 3.2x300mm | Sigma | GE29-0915-98 |
| Ultrathin carbon film on lacey carbon support 400M Cu | Ted Pella | 01824 |
| Quantifoil R 1.2/1.3 Au 300 mesh grids | Quantifoil | |
| Tecnai F20 | FEI | |
| Leica EM GP | Leica | |
| Titan Krios | FEI | |
| Pierce Fab Preparation Kit | Thermo Scientific | 44985 |
| Amicon Ultra-4 Centrifugal Filter Units 100kDa | Millipore | UFC810024 |
| Amicon Ultra-4 Centrifugal Filter Units 10kDa | Millipore | UFC801024 |
| QIAquick Gel Extraction Kit | Qiagen | 28706 |
| QIAquick Nucleotide Removal Kit | Qiagen | 28304 |
| QIAprep Spin Miniprep Kit | Qiagen | 27106 |
| QIAGEN Large-Construct Kit | Qiagen | 12462 |
| Immobilon-P | Merck Millipore Ltd. | IPVH00010 |
| Optical bottom 96-well black sided culture plates | Thermo Scientific | 165305 |
| FACSCalibur flow cytometer | Becton Dickenson | |
| Synergy H1 Multi-mode Reader | Biotek | |
| Nunclon Delta Surface 12-well plates | Thermo Scientific | 150628 |
| Cell Culture 6-well plates | Corning | 3506 |
| Microscope cover glass 18mm No. 1 | Fisher Scientific | 12-545-100 |
| Orbitrap Fusion mass Spectrometer | Thermo Scientific | |
| Acquity M-Class liquid chromatograph | Waters Corporation | |
| 1.8 micron C18 stationary phase beads | Dr. Maisch | http://www.dr-maisch.com |

**Supporting Information References.**

1. Heldwein EE, Lou H, Bender FC, Cohen GH, Eisenberg RJ, Harrison SC. Crystal structure of glycoprotein B from herpes simplex virus 1. Science. 2006;313(5784):217-20. doi: 10.1126/science.1126548. PubMed PMID: 16840698.
2. Vallbracht M, Brun D, Tassinari M, Vaney MC, Pehau-Arnaudet G, Guardado-Calvo P, et al. Structure-function dissection of the Pseudorabies virus glycoprotein B fusion loops. J Virol. 2017. doi: 10.1128/JVI.01203-17. PubMed PMID: 29046441; PubMed Central PMCID: PMCPMC5730762.
3. Burke HG, Heldwein EE. Crystal Structure of the Human Cytomegalovirus Glycoprotein B. PLoS Pathog. 2015;11(10):e1005227. doi: 10.1371/journal.ppat.1005227. PubMed PMID: 26484870; PubMed Central PMCID: PMCPMC4617298.
4. Backovic M, Longnecker R, Jardetzky TS. Structure of a trimeric variant of the Epstein-Barr virus glycoprotein B. Proc Natl Acad Sci U S A. 2009;106(8):2880-5. doi: 10.1073/pnas.0810530106. PubMed PMID: 19196955; PubMed Central PMCID: PMCPMC2650359.
5. Vollmer B, Prazak V, Vasishtan D, Jefferys EE, Hernandez-Duran A, Vallbracht M, et al. The prefusion structure of herpes simplex virus glycoprotein B. Sci Adv. 2020;6(39). doi: 10.1126/sciadv.abc1726. PubMed PMID: 32978151; PubMed Central PMCID: PMCPMC7518877.
6. van Genderen IL, Brandimarti R, Torrisi MR, Campadelli G, van Meer G. The phospholipid composition of extracellular herpes simplex virions differs from that of host cell nuclei. Virology. 1994;200(2):831-6. doi: 10.1006/viro.1994.1252. PubMed PMID: 8178468.
7. Oliver SL, Sommer M, Zerboni L, Rajamani J, Grose C, Arvin AM. Mutagenesis of varicella-zoster virus glycoprotein B: putative fusion loop residues are essential for viral replication, and the furin cleavage motif contributes to pathogenesis in skin tissue in vivo. J Virol. 2009;83(15):7495-506. doi: 10.1128/JVI.00400-09. PubMed PMID: 19474103; PubMed Central PMCID: PMCPMC2708640.
8. Tischer BK, Kaufer BB, Sommer M, Wussow F, Arvin AM, Osterrieder N. A self-excisable infectious bacterial artificial chromosome clone of varicella-zoster virus allows analysis of the essential tegument protein encoded by ORF9. J Virol. 2007;81(23):13200-8. doi: 10.1128/JVI.01148-07. PubMed PMID: 17913822; PubMed Central PMCID: PMCPMC2169085.
9. Tischer BK, Smith GA, Osterrieder N. En passant mutagenesis: a two step markerless red recombination system. Methods Mol Biol. 2010;634:421-30. doi: 10.1007/978-1-60761-652-8_30. PubMed PMID: 20677001.
10. Yang E, Arvin AM, Oliver SL. The cytoplasmic domain of varicella-zoster virus glycoprotein H regulates syncytia formation and skin pathogenesis. PLoS Pathog. 2014;10(5):e1004173. doi: 10.1371/journal.ppat.1004173. PubMed PMID: 24874654; PubMed Central PMCID: PMCPMC4038623.
11. Yang E, Arvin AM, Oliver SL. Role for the alphaV Integrin Subunit in Varicella-Zoster Virus-Mediated Fusion and Infection. J Virol. 2016;90(16):7567-78. doi: 10.1128/JVI.00792-16. PubMed PMID: 27279620; PubMed Central PMCID: PMCPMC4984616.
12. Oliver SL, Xing Y, Chen DH, Roh SH, Pintilie GD, Bushnell DA, et al. A glycoprotein B-neutralizing antibody structure at 2.8 A uncovers a critical domain for herpesvirus fusion

initiation. Nat Commun. 2020;11(1):4141. doi: 10.1038/s41467-020-17911-0. PubMed PMID: 32811830; PubMed Central PMCID: PMCPMC7435202.
