## Supplementary figures and images for "The N-terminus of varicella-zoster virus glycoprotein B has a functional role in fusion"

### S1A Fig

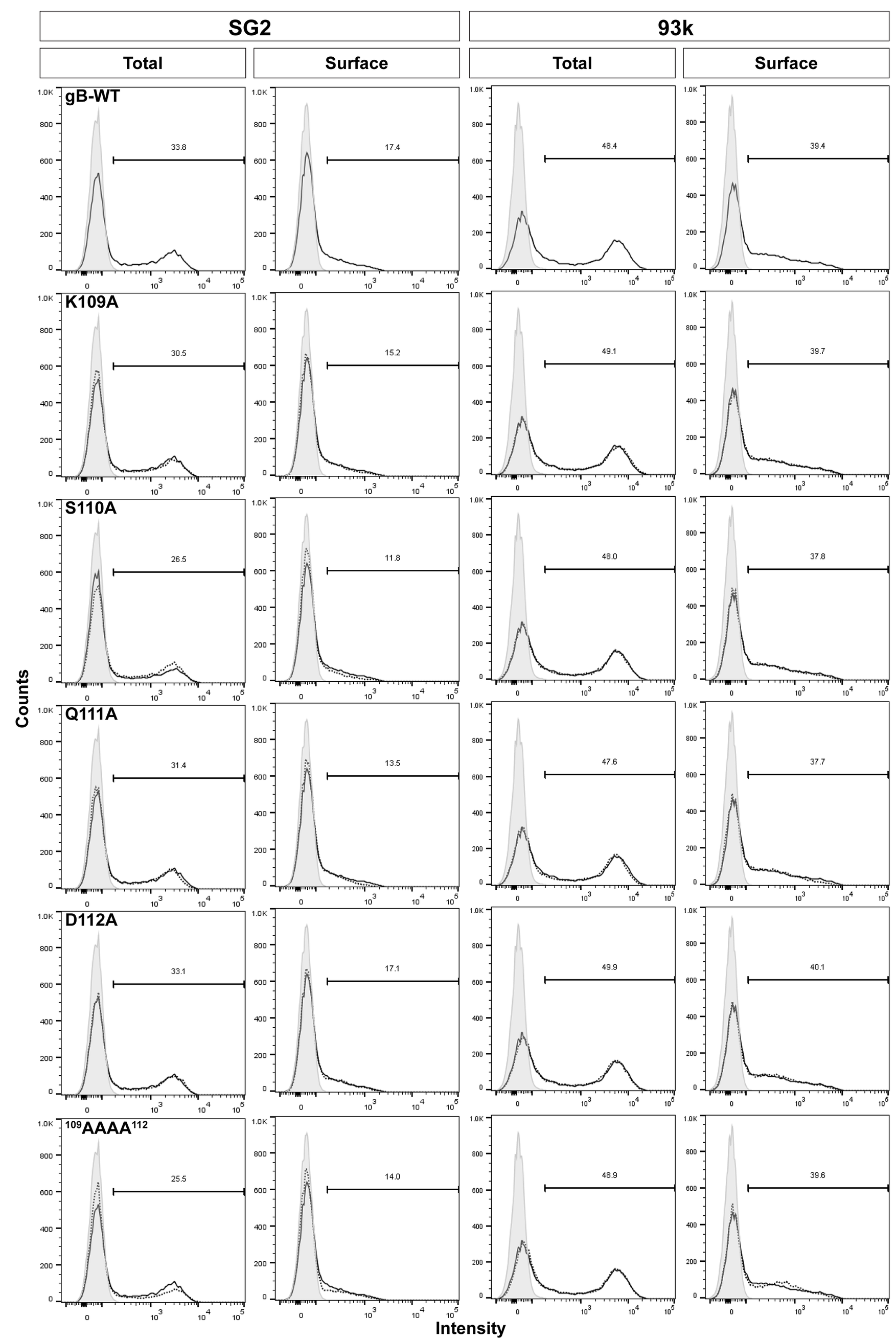

### S1B Fig

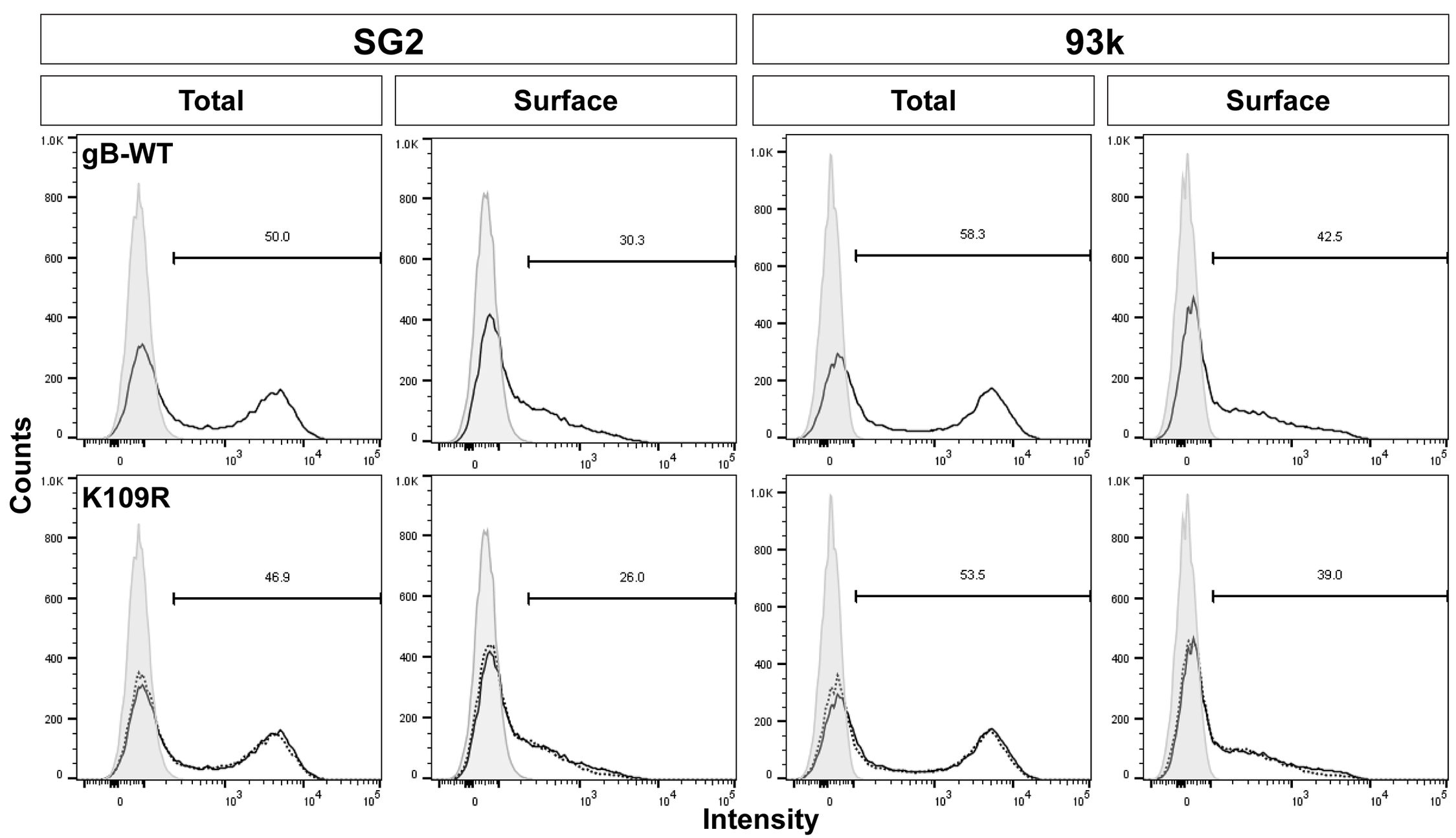

### S2 Fig

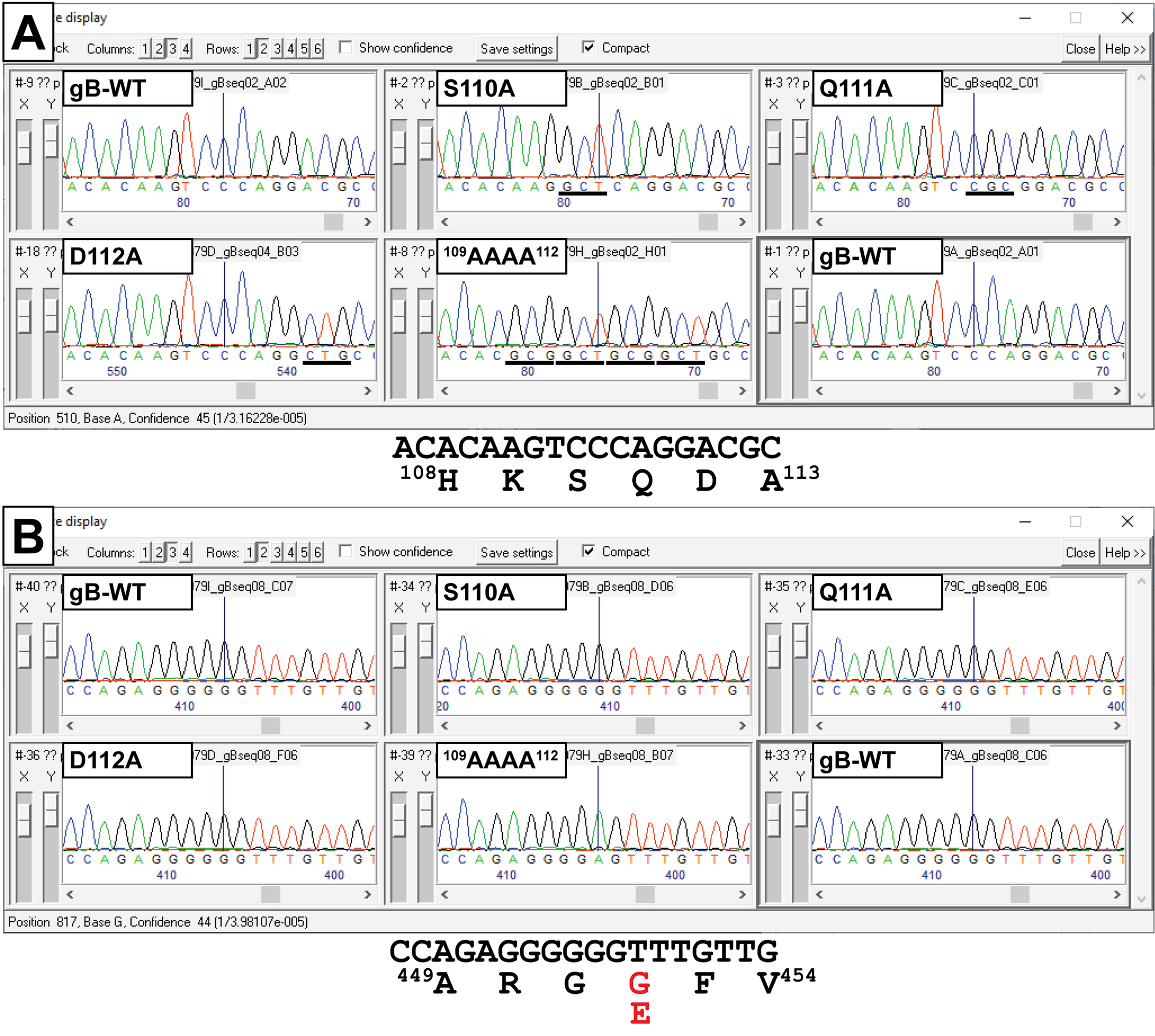

### S3 Fig

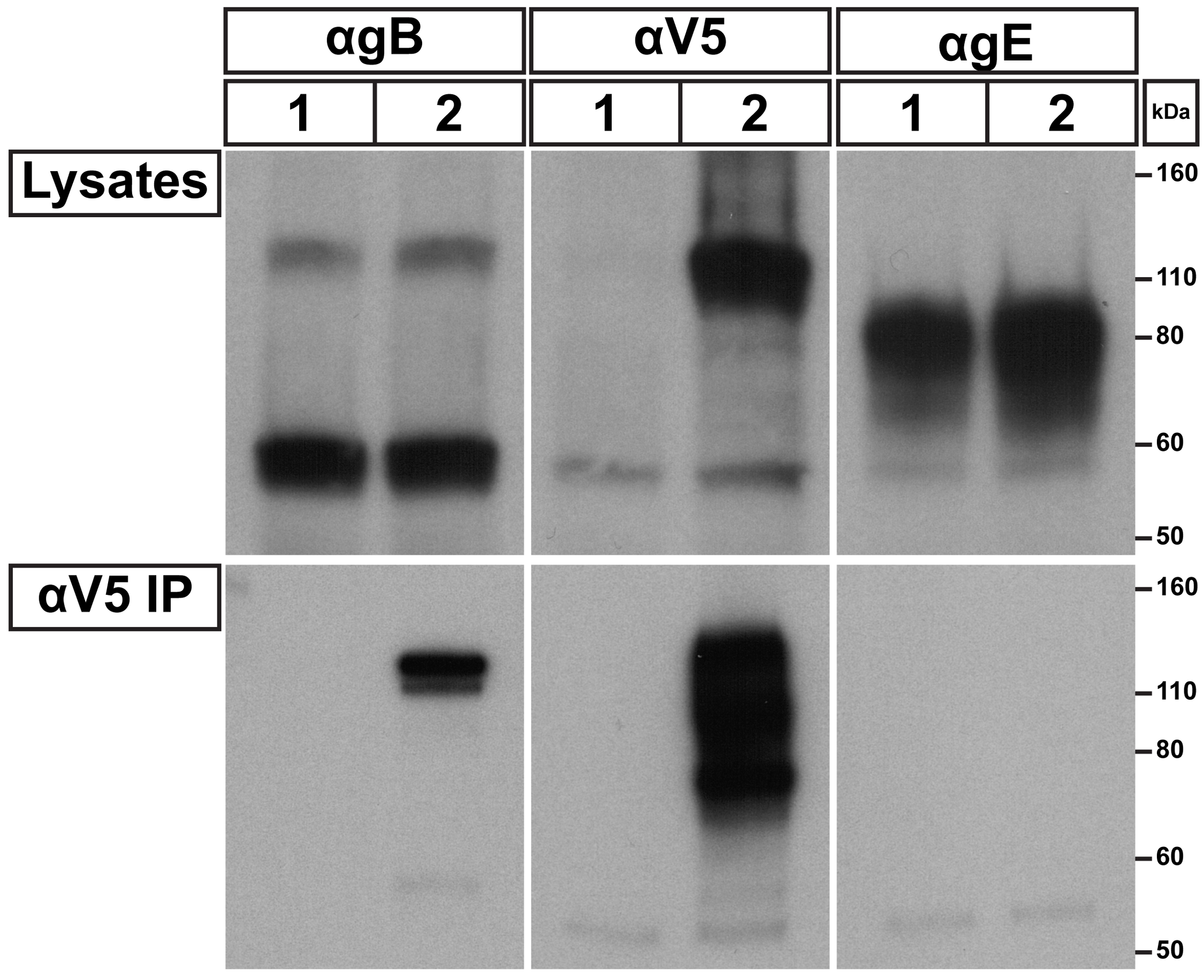
