## Supplementary material for "The N-terminus of varicella-zoster virus glycoprotein B has a functional role in fusion": S2 Validation Report

### Full wwPDB EM Map Validation Report (i)

Aug 31, 2020 – 11:57 AM EDT

| | | |
|---|---|---|
| EMDB ID | : | EMD-22520 |
| Title | : | The N-terminus of varicella-zoster virus glycoprotein B has a functional role in fusion |
| Deposited on | : | 2020-08-27 |
| Resolution | : | 9.00 Å(reported) |

This is a Full wwPDB EM Map Validation Report.

This report is produced by the wwPDB biocuration pipeline after annotation of the structure.

We welcome your comments at **
A user guide is available at
https://www.wwpdb.org/validation/2017/EMMapValidationReportHelp
with specific help available everywhere you see the (i) symbol.

CONFIDENTIAL VALIDATION REPORT

The following versions of software and data (see references (i)) were used in the production of this report:

| | | |
|---|---|---|
| EMDB validation analysis | : | 0.0.0.dev33 |
| Validation Pipeline (wwPDB-VP) | : | 2.13.1 |

### 1 Experimental information (i)

| Property | Value | Source |
|---|---|---|
| EM reconstruction method | SINGLE PARTICLE | Depositor |
| Imposed symmetry | POINT, Not provided | Depositor |
| Number of particles used | 59610 | Depositor |
| Resolution determination method | FSC 0.143 CUT-OFF | Depositor |
| CTF correction method | NONE | Depositor |
| Microscope | FEI TITAN KRIOS | Depositor |
| Voltage (kV) | 300 | Depositor |
| Electron dose $(e^-/\text{Å}^2)$ | 7.8 | Depositor |
| Minimum defocus (nm) | Not provided | Depositor |
| Maximum defocus (nm) | Not provided | Depositor |
| Magnification | Not provided | Depositor |
| Image detector | GATAN K2 SUMMIT (4k x 4k) | Depositor |
| Maximum map value | 11.432 | Depositor |
| Minimum map value | -7.053 | Depositor |
| Average map value | 0.000 | Depositor |
| Map value standard deviation | 0.273 | Depositor |
| Recommended contour level | 2.16 | Depositor |
| Map size (Å) | 513.20026, 513.20026, 513.20026 | Depositor |
| Map dimensions | 200, 200, 200 | Depositor |
| Map angles (°) | 90.0, 90.0, 90.0 | Depositor |
| Pixel spacing (Å) | 2.5660012, 2.5660012, 2.5660012 | Depositor |

CONFIDENTIAL VALIDATION REPORT

### 2 Map visualisation (i)

This section contains visualisations of the EMDB entry EMD-22520. These are intended to permit visual inspection of the internal detail of the map and identification of artifacts.

#### 2.1 Orthogonal projections (i)

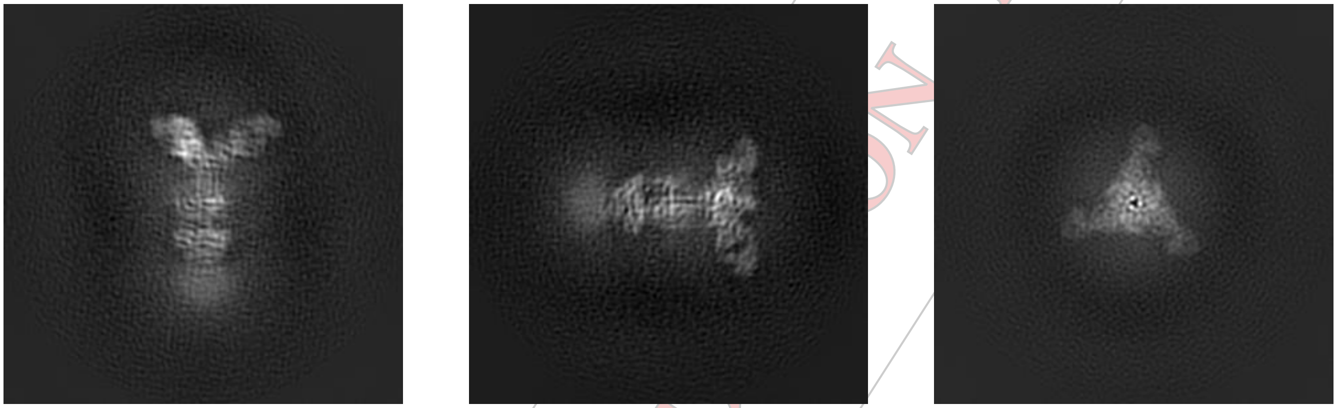

X Y Z

The images above show the map projected in three orthogonal projections, in greyscale.

#### 2.2 Central slices (i)

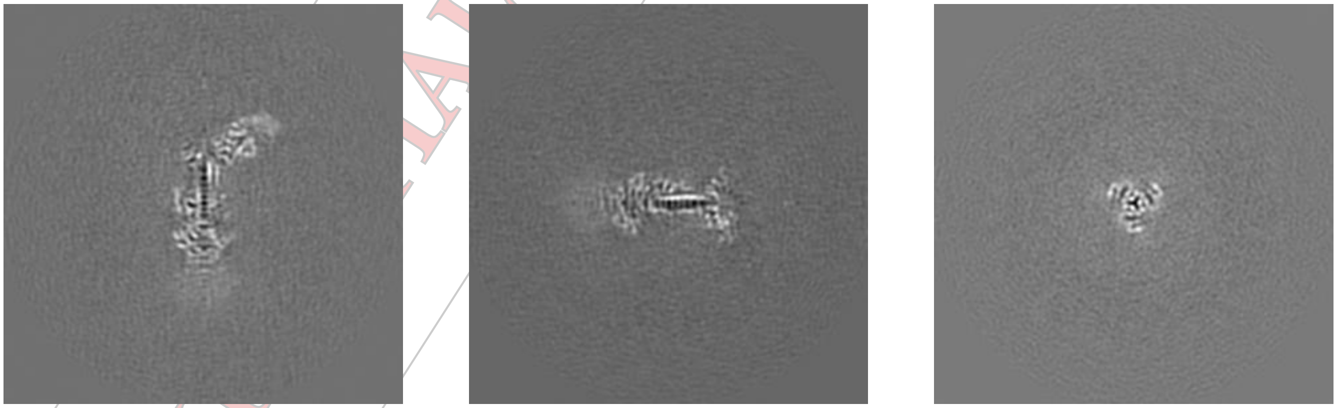

X Y Z

The images above show central slices of the map in three orthogonal directions, in greyscale.

CONFIDENTIAL VALIDATION REPORT

#### 2.3 Largest variance slices (i)

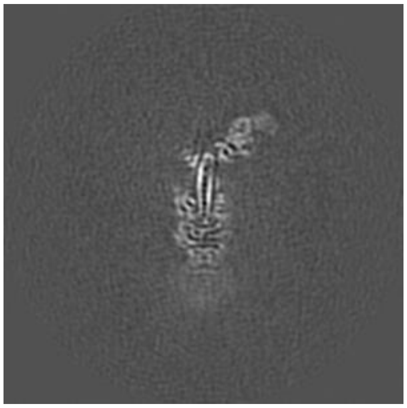
X Index: 98

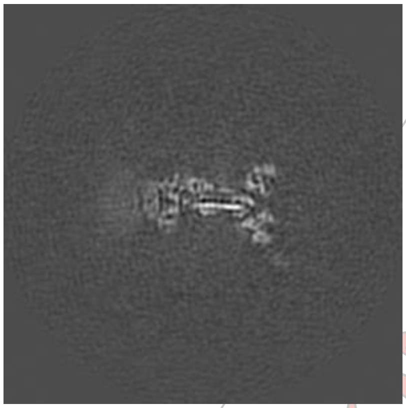
Y Index: 98

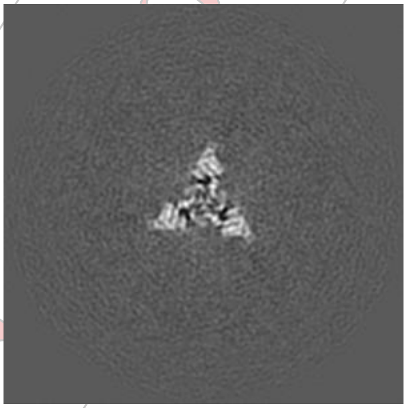
Z Index: 125

The images above show the highest variance slices of the map in three orthogonal directions, in greyscale.

#### 2.4 Orthogonal surface views (i)

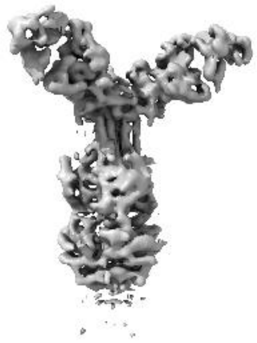
X

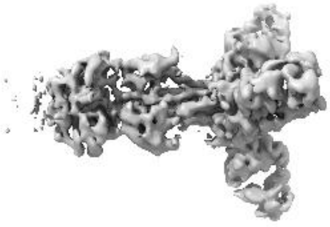
Y

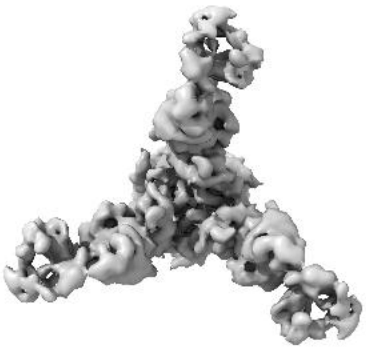
Z

The images above show the 3D surface view of the map at the recommended contour level 2.16. This in conjunction with the slice images can indicate whether an appropriate contour level has been selected.

#### 2.5 Mask visualisation (i)

This section was not generated. No masks were provided.

CONFIDENTIAL - VALIDATION REPORT

### 3 Map analysis (i)

This section contains the results of statistical analysis of the map.

#### 3.1 Map-value distribution (i)

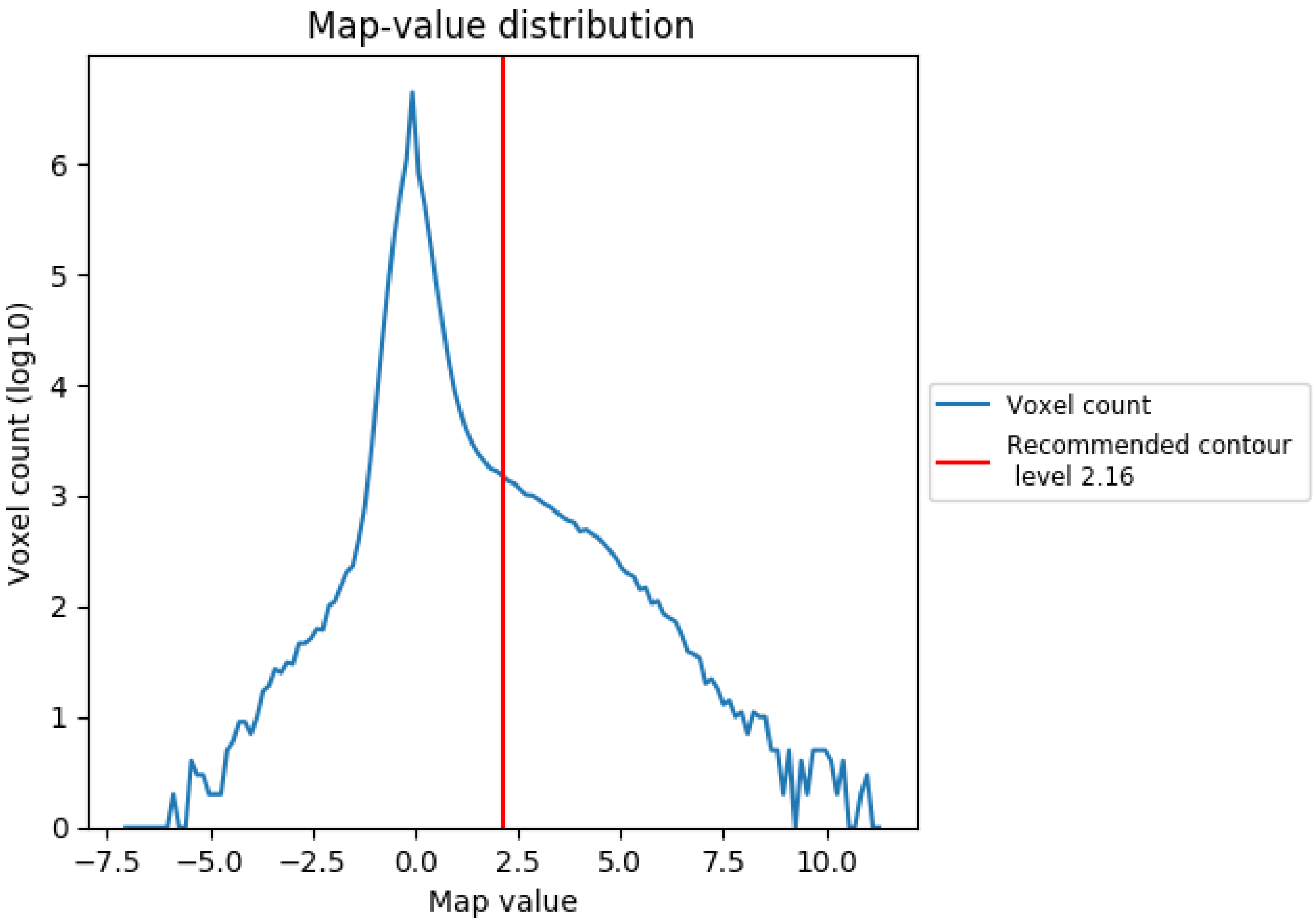


The map-value distribution is plotted in 128 intervals along the x-axis. The y-axis is logarithmic. A spike in this graph at zero usually indicates that the volume has been masked.

CONFIDENTIAL REPORT

#### 3.2 Volume estimate (i)

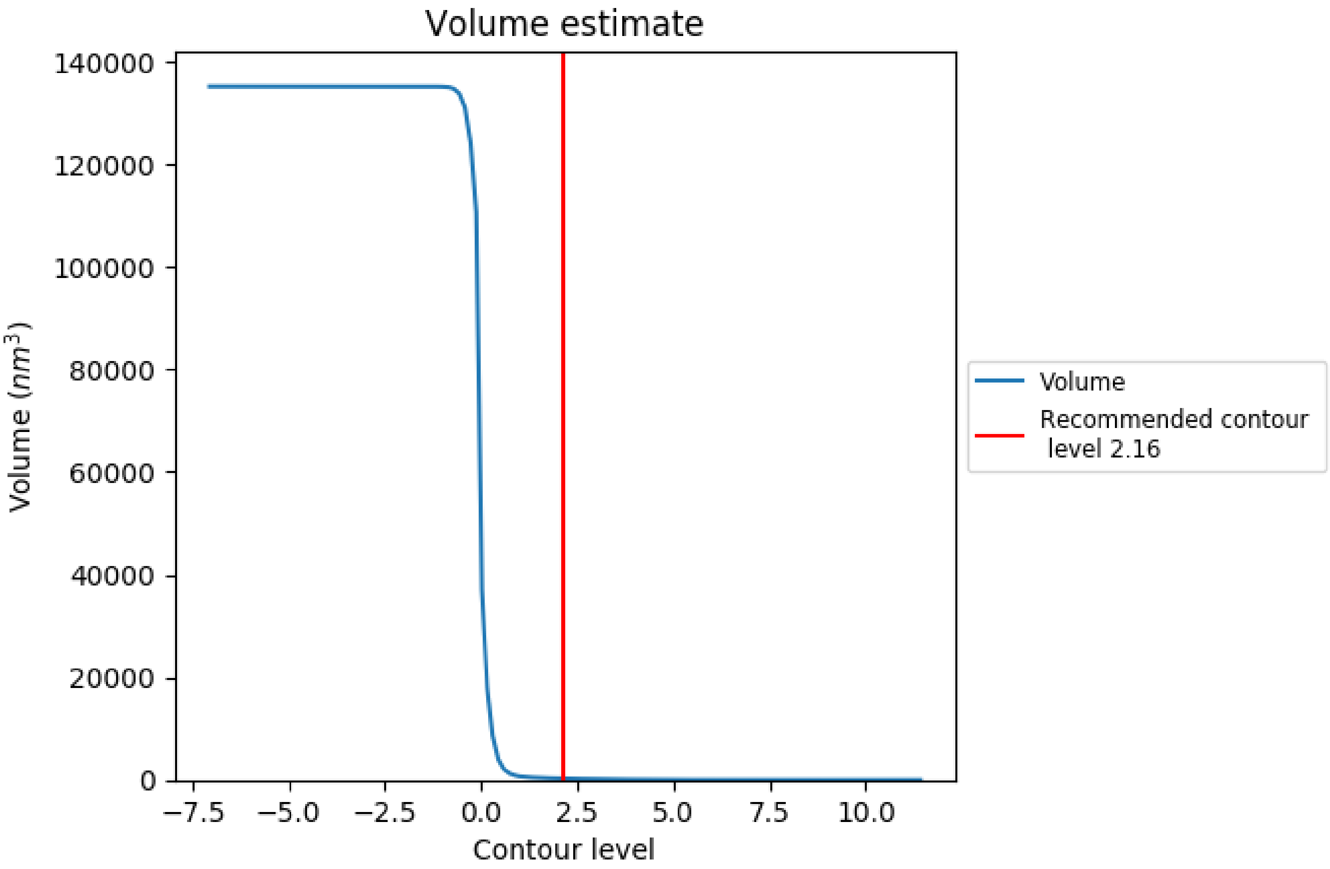


The volume at the recommended contour level is 277 $nm^3$; this corresponds to an approximate mass of 250 kDa.

The volume estimate graph shows how the enclosed volume varies with the contour level. The recommended contour level is shown as a vertical line and the intersection between the line and the curve gives the volume of the enclosed surface at the given level.

CONFIDENTIAL REPORT

#### 3.3 Rotationally averaged power spectrum (i)

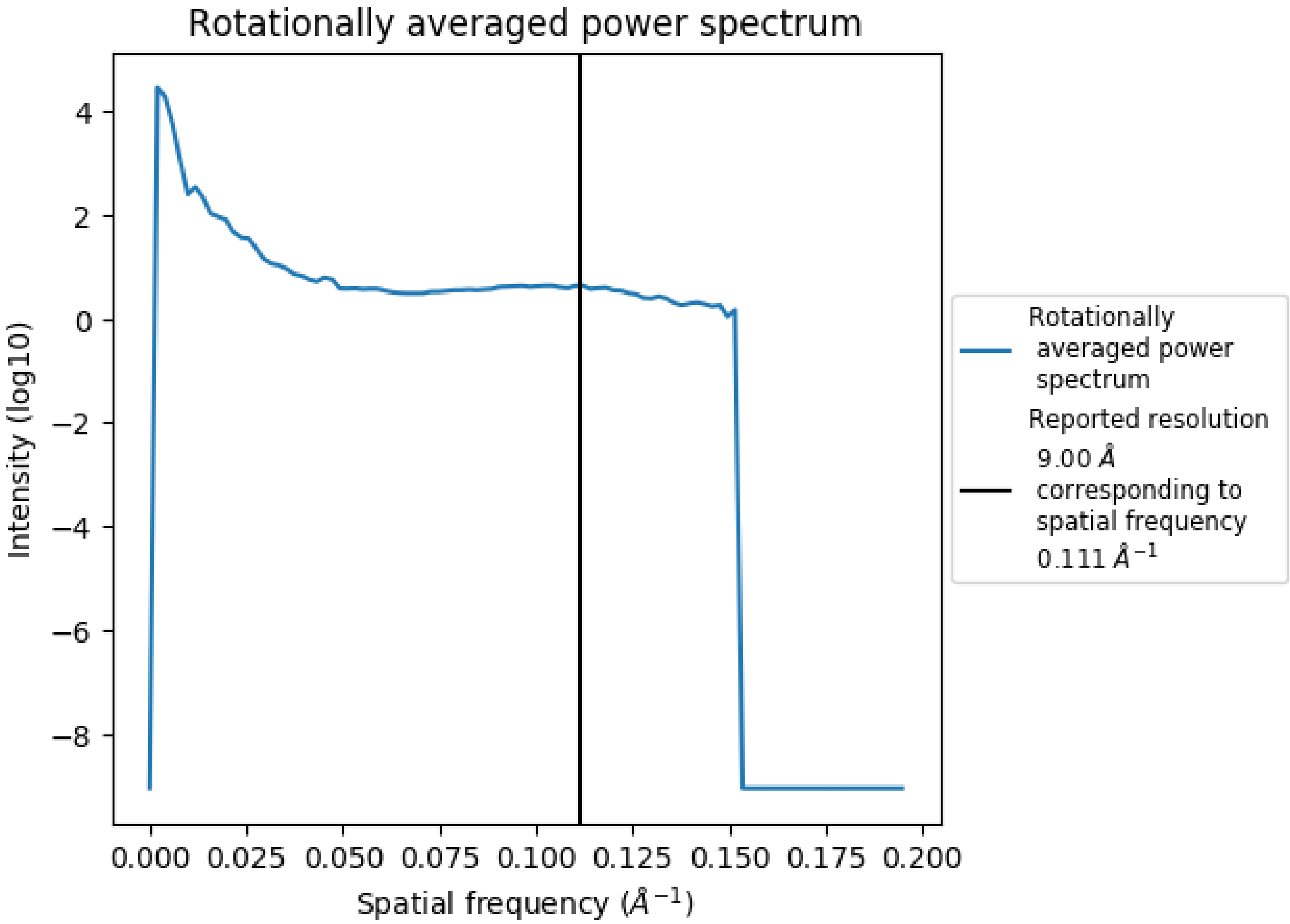


CONFIDENTIAL REPORT

### 4 Fourier-Shell correlation (i)

This section was not generated. No FSC curve or half maps provided.

CONFIDENTIAL VALIDATION REPORT
