## Supplementary material for "The N-terminus of varicella-zoster virus glycoprotein B has a functional role in fusion": S3 Validation Report

### Preliminary Full wwPDB EM Map/Model Validation Report (i)

Sep 8, 2020 – 11:02 AM EDT

Deposition ID : D_1000251554

This is a Preliminary Full wwPDB EM Map/Model Validation Report.

This report is produced by the wwPDB Deposition System during initial deposition but before annotation of the structure.

We welcome your comments at **
A user guide is available at
https://www.wwpdb.org/validation/2017/EMValidationReportHelp
with specific help available everywhere you see the (i) symbol.

PRELIMINARY VALIDATION REPORT

The following versions of software and data (see references (i)) were used in the production of this report:

| | | |
|---|---|---|
| EMDB validation analysis | : | 0.0.0.dev33 |
| Mogul | : | 1.8.5 (274361), CSD as541be (2020) |
| MolProbity | : | 4.02b-467 |
| Percentile statistics | : | 20191225.v01 (using entries in the PDB archive December 25th 2019) |
| Ideal geometry (proteins) | : | Engh & Huber (2001) |
| Ideal geometry (DNA, RNA) | : | Parkinson et al. (1996) |
| Validation Pipeline (wwPDB-VP) | : | 2.14.2 |

### 1 Overall quality at a glance (i)

The following experimental techniques were used to determine the structure:
*ELECTRON MICROSCOPY*

The reported resolution of this entry is 3.90 Å.

Percentile scores (ranging between 0-100) for global validation metrics of the entry are shown in the following graphic. The table shows the number of entries on which the scores are based.

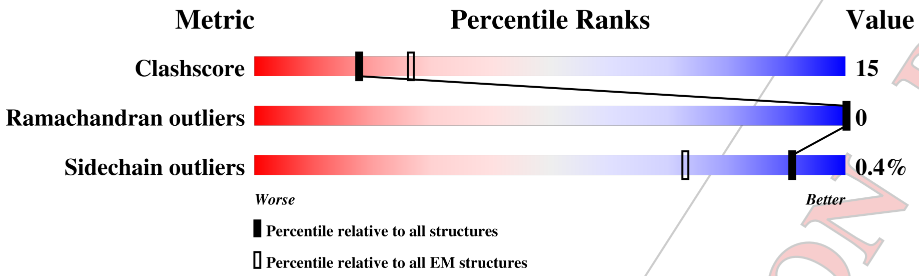


| Metric | Whole archive (#Entries) | EM structures (#Entries) |
|---|---|---|
| Clashscore | 158937 | 4297 |
| Ramachandran outliers | 154571 | 4023 |
| Sidechain outliers | 154315 | 3826 |

The table below summarises the geometric issues observed across the polymeric chains and their fit to the map. The red, orange, yellow and green segments on the bar indicate the fraction of residues that contain outliers for >=3, 2, 1 and 0 types of geometric quality criteria respectively. A grey segment represents the fraction of residues that are not modelled. The numeric value for each fraction is indicated below the corresponding segment, with a dot representing fractions <=5% The upper red bar (where present) indicates the fraction of residues that have poor fit to the EM map (all atom inclusion < 40%). The numeric value is given above the bar.

| Mol | Chain | Length | Quality of chain |
|---|---|---|---|
| 1 | A | 931 | • 45% 17% 37% |
| 1 | B | 931 | • 45% 17% 37% |
| 1 | C | 931 | • 45% 17% 37% |
| 2 | D | 6 | 17% 33% 50% 17% |
| 2 | E | 6 | 17% 33% 33% 33% |
| 2 | F | 6 | 17% 33% 50% 17% |

PRELIMINARY VALIDATION REPORT

#### 2 Entry composition (i)

There are 3 unique types of molecules in this entry. The entry contains 14409 atoms, of which 0 are hydrogens and 0 are deuteriums.

In the tables below, the AltConf column contains the number of residues with at least one atom in alternate conformation and the Trace column contains the number of residues modelled with at most 2 atoms.

- Molecule 1 is a protein called Varicella-zoster virus glycoprotein B.

| Mol | Chain | Residues | Atoms | | | | | AltConf | Trace |
|---|---|---|---|---|---|---|---|---|---|
| | | | Total | C | N | O | S | | |
| 1 | A | 584 | 4717 | 2981 | 823 | 891 | 22 | 0 | 0 |
| 1 | B | 584 | 4717 | 2981 | 823 | 891 | 22 | 0 | 0 |
| 1 | C | 584 | 4717 | 2981 | 823 | 891 | 22 | 0 | 0 |

- Molecule 2 is an oligosaccharide called alpha-D-mannopyranose-(1-3)-alpha-D-mannopyranose-(1-6)-[alpha-D-mannopyranose-(1-3)]alpha-D-mannopyranose-(1-4)-2-acetamido-2-deoxy-beta-D-glucopyranose-(1-4)-2-acetamido-2-deoxy-beta-D-glucopyranose.

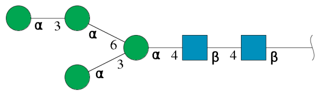

| Mol | Chain | Residues | Atoms | | | | AltConf | Trace |
|---|---|---|---|---|---|---|---|---|
| | | | Total | C | N | O | | |
| 2 | D | 6 | 72 | 40 | 2 | 30 | 0 | 0 |
| 2 | E | 6 | 72 | 40 | 2 | 30 | 0 | 0 |
| 2 | F | 6 | 72 | 40 | 2 | 30 | 0 | 0 |

- Molecule 3 is 2-acetamido-2-deoxy-beta-D-glucopyranose (three-letter code: NAG) (formula: $C_8H_{15}NO_6$).

PRELIMINARY VALIDATION REPORT

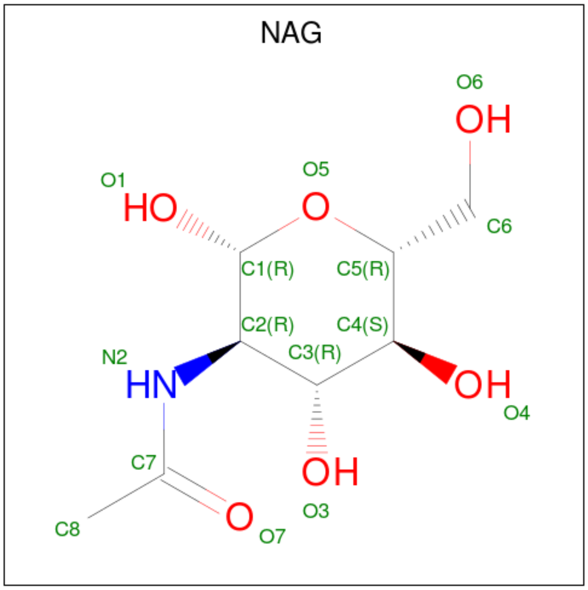


| Mol | Chain | Residues | Atoms | AltConf |
|---|---|---|---|---|
| 3 | A | 1 | Total C N O<br>14 8 1 5 | 0 |
| 3 | B | 1 | Total C N O<br>14 8 1 5 | 0 |
| 3 | C | 1 | Total C N O<br>14 8 1 5 | 0 |

PRELIMINARY VALIDATION REPORT

### 3 Residue-property plots ⓘ

These plots are drawn for all protein, RNA, DNA and oligosaccharide chains in the entry. The first graphic for a chain summarises the proportions of the various outlier classes displayed in the second graphic. The second graphic shows the sequence view annotated by issues in geometry and atom inclusion in map density. Residues are color-coded according to the number of geometric quality criteria for which they contain at least one outlier: green = 0, yellow = 1, orange = 2 and red = 3 or more. A red diamond above a residue indicates a poor fit to the EM map for this residue (all atom inclusion < 40%). Stretches of 2 or more consecutive residues without any outlier are shown as a green connector. Residues present in the sample, but not in the model, are shown in grey.

- Molecule 1: Varicella-zoster virus glycoprotein B

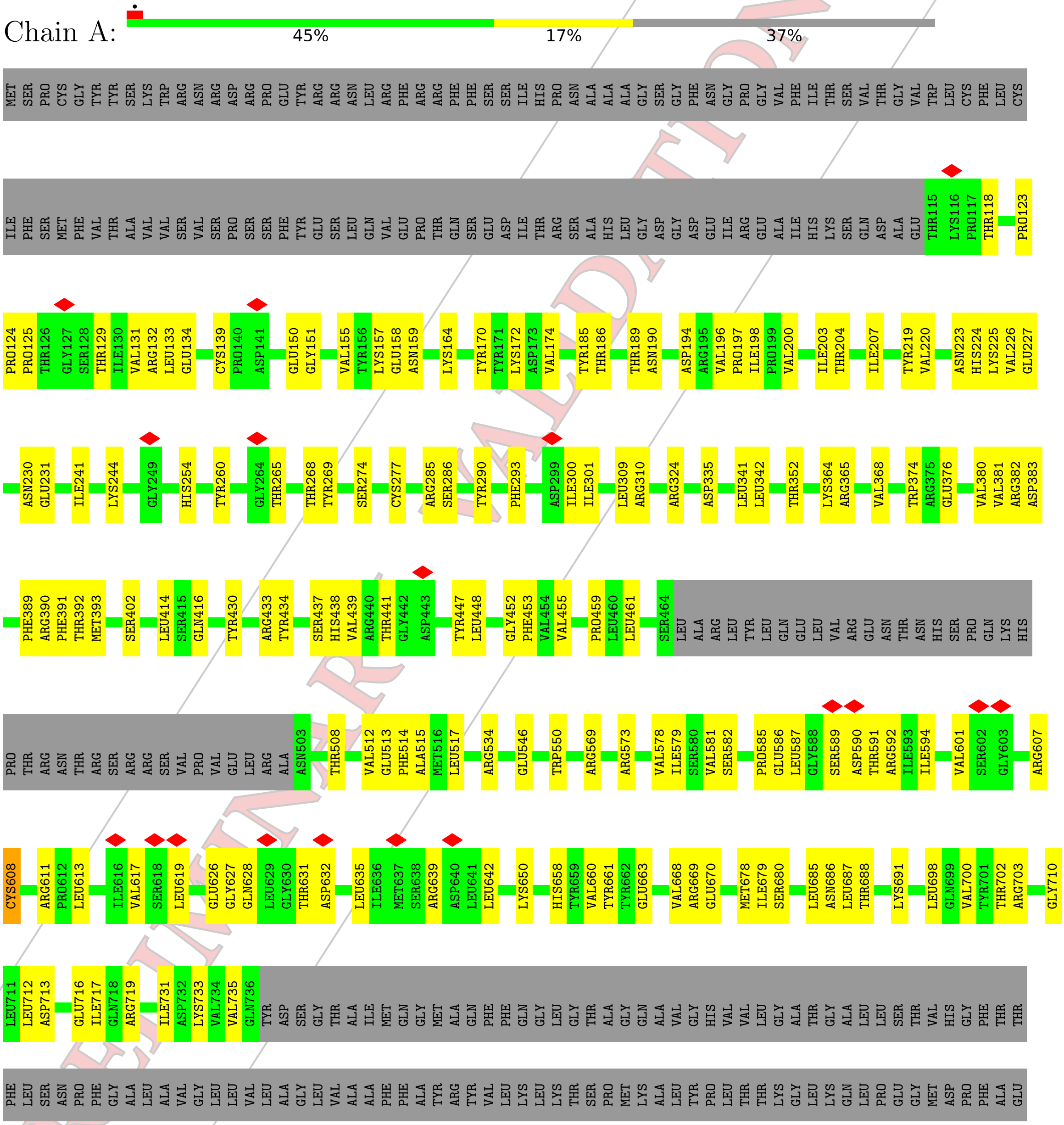


PRELIMINARY VALIDATION REPORT

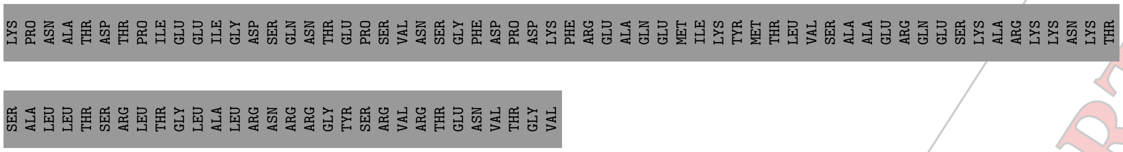


• Molecule 1: Varicella-zoster virus glycoprotein B

Chain B:
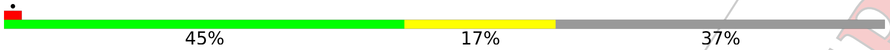


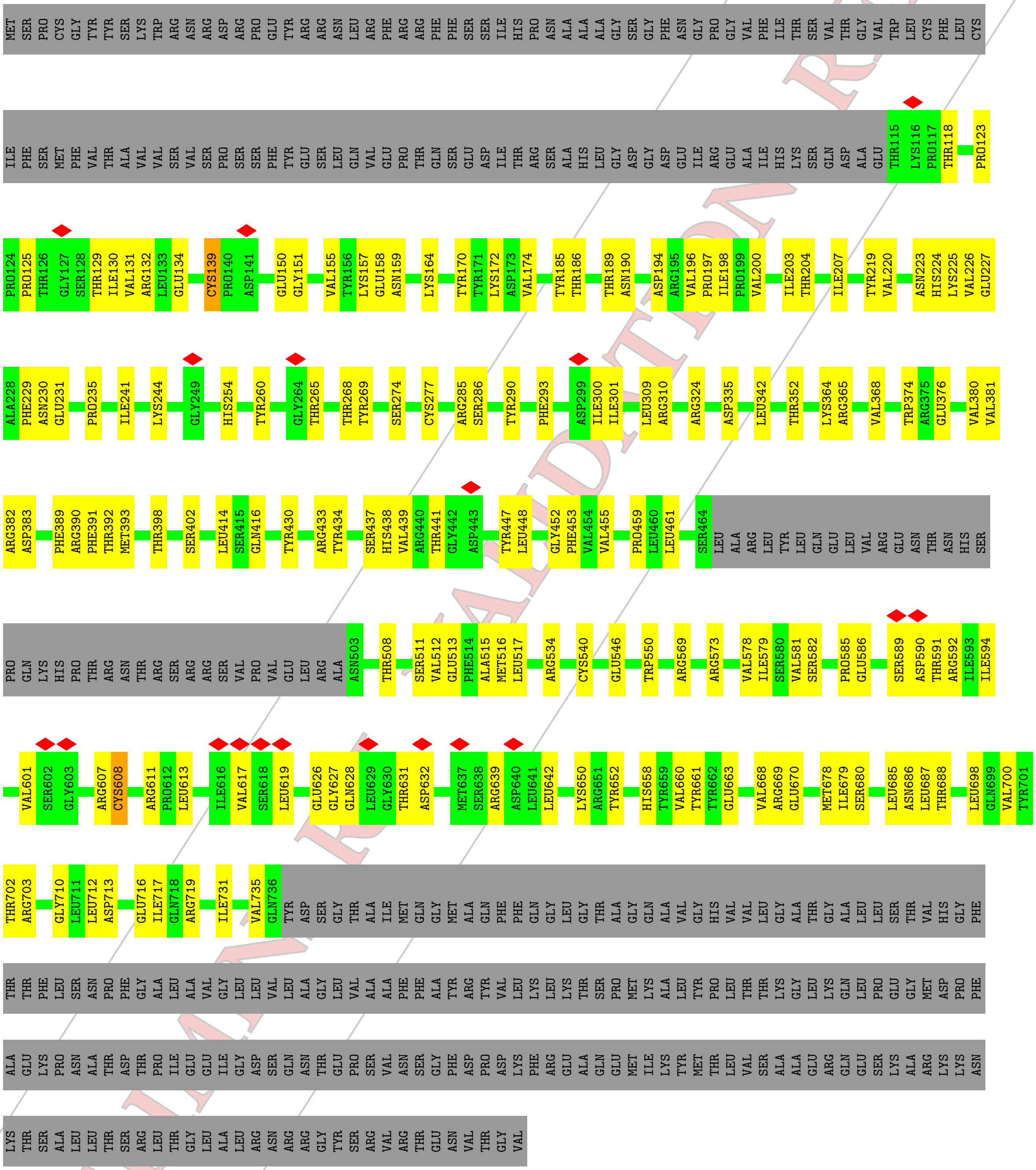


• Molecule 1: Varicella-zoster virus glycoprotein B

Chain C:
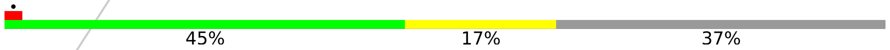


PRELIMINARY VALIDATION REPORT

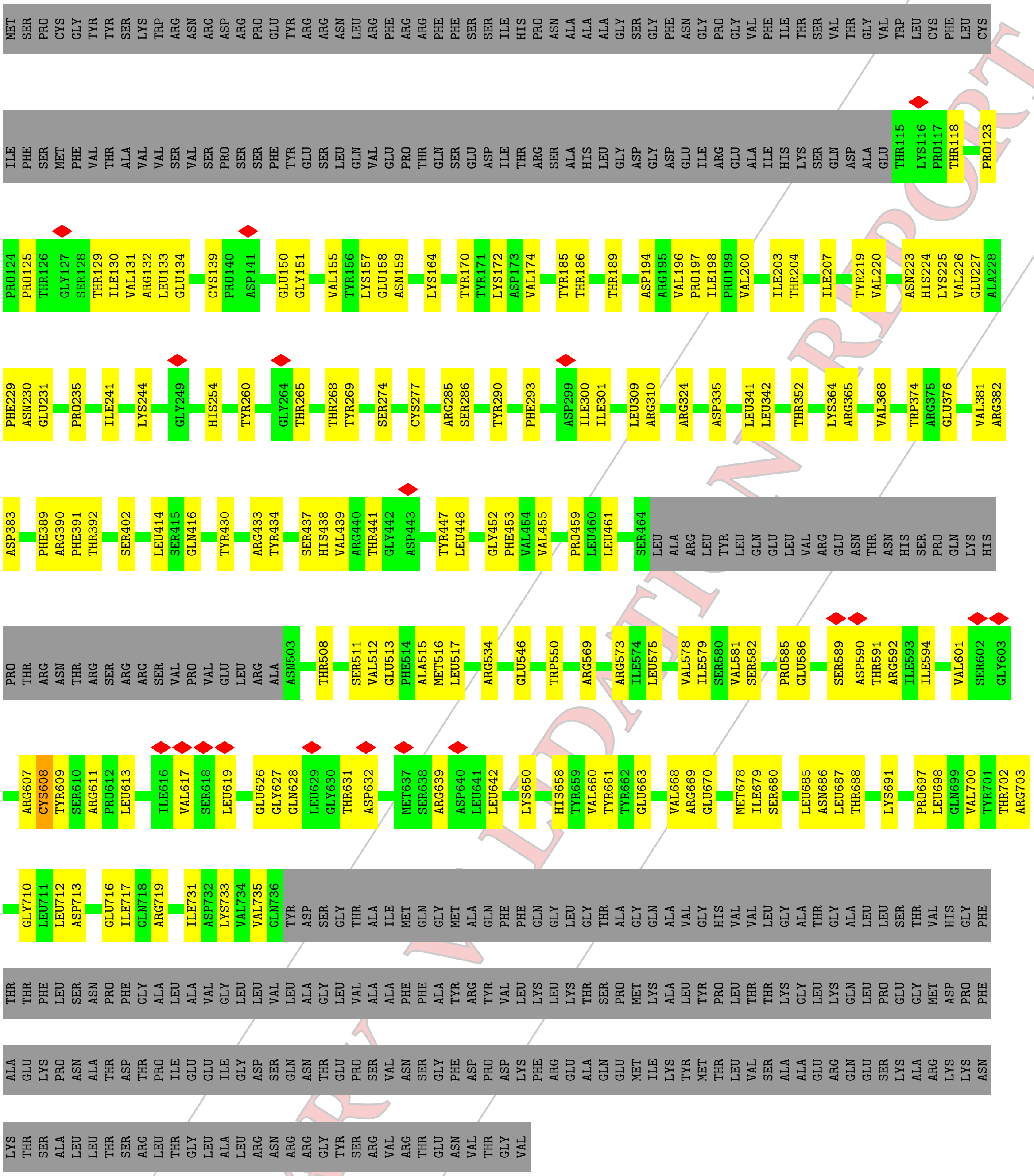

• Molecule 2: alpha-D-mannopyranose-(1-3)-alpha-D-mannopyranose-(1-6)-[alpha-D-mannopyranose-(1-3)]alpha-D-mannopyranose-(1-4)-2-acetamido-2-deoxy-beta-D-glucopyranose-(1-4)-2-acetamido-2-deoxy-beta-D-glucopyranose

Chain D: 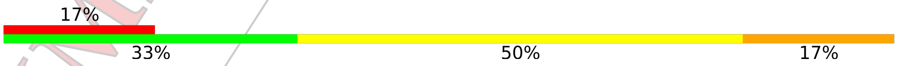


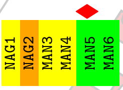


• Molecule 2: alpha-D-mannopyranose-(1-3)-alpha-D-mannopyranose-(1-6)-[alpha-D-mannopyranose-(1-3)]alpha-D-mannopyranose-(1-4)-2-acetamido-2-deoxy-beta-D-glucopyranose-(1-4)-2-aceta

PRELIMINARY VALIDATION REPORT

mido-2-deoxy-beta-D-glucopyranose

Chain E: 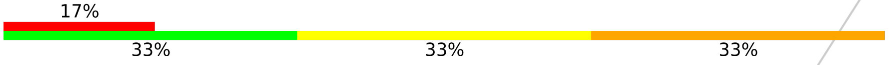


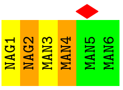


• Molecule 2: alpha-D-mannopyranose-(1-3)-alpha-D-mannopyranose-(1-6)-[alpha-D-mannopyranose-(1-3)]alpha-D-mannopyranose-(1-4)-2-acetamido-2-deoxy-beta-D-glucopyranose-(1-4)-2-acetamido-2-deoxy-beta-D-glucopyranose

Chain F: 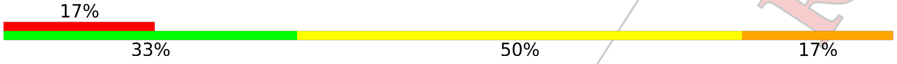


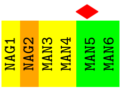


PRELIMINARY VALIDATION REPORT

### 4 Experimental information ⓘ

| Property | Value | Source |
|---|---|---|
| EM reconstruction method | Not provided | Depositor |
| Imposed symmetry | POINT, Not provided | Depositor |
| Number of images used | Not provided | Depositor |
| Resolution determination method | Not provided | Depositor |
| CTF correction method | Not provided | Depositor |
| Microscope | Not provided | Depositor |
| Voltage (kV) | Not provided | Depositor |
| Electron dose ($e^{-}/\text{Å}^2$) | Not provided | Depositor |
| Minimum defocus (nm) | Not provided | Depositor |
| Maximum defocus (nm) | Not provided | Depositor |
| Magnification | Not provided | Depositor |
| Image detector | Not provided | Depositor |
| Maximum map value | 0.148 | Depositor |
| Minimum map value | -0.083 | Depositor |
| Average map value | 0.000 | Depositor |
| Map value standard deviation | 0.002 | Depositor |
| Recommended contour level | 0.02 | Depositor |
| Map size (Å) | 423.99997, 423.99997, 423.99997 | Depositor |
| Map dimensions | 400, 400, 400 | Depositor |
| Map angles (°) | 90.0, 90.0, 90.0 | Depositor |
| Pixel spacing (Å) | 1.06, 1.06, 1.06 | Depositor |

PRELIMINARY VALIDATION REPORT

### 5 Model quality (i)

#### 5.1 Standard geometry (i)

Bond lengths and bond angles in the following residue types are not validated in this section: NAG, MAN

The Z score for a bond length (or angle) is the number of standard deviations the observed value is removed from the expected value. A bond length (or angle) with $|Z| > 5$ is considered an outlier worth inspection. RMSZ is the root-mean-square of all Z scores of the bond lengths (or angles).

| Mol | Chain | Bond lengths | | Bond angles | |
|---|---|---|---|---|---|
| | | RMSZ | $\#\|Z\| > 5$ | RMSZ | $\#\|Z\| > 5$ |
| 1 | A | 0.36 | 0/4827 | 0.54 | 0/6555 |
| 1 | B | 0.36 | 0/4827 | 0.54 | 0/6555 |
| 1 | C | 0.36 | 0/4827 | 0.54 | 0/6555 |
| All | All | 0.36 | 0/14481 | 0.54 | 0/19665 |

There are no bond length outliers.

There are no bond angle outliers.

There are no chirality outliers.

There are no planarity outliers.

#### 5.2 Too-close contacts (i)

In the following table, the Non-H and H(model) columns list the number of non-hydrogen atoms and hydrogen atoms in the chain respectively. The H(added) column lists the number of hydrogen atoms added and optimized by MolProbity. The Clashes column lists the number of clashes within the asymmetric unit, whereas Symm-Clashes lists symmetry related clashes.

| Mol | Chain | Non-H | H(model) | H(added) | Clashes | Symm-Clashes |
|---|---|---|---|---|---|---|
| 1 | A | 4717 | 0 | 4597 | 177 | 0 |
| 1 | B | 4717 | 0 | 4597 | 188 | 0 |
| 1 | C | 4717 | 0 | 4597 | 177 | 0 |
| 2 | D | 72 | 0 | 61 | 8 | 0 |
| 2 | E | 72 | 0 | 61 | 7 | 0 |
| 2 | F | 72 | 0 | 61 | 8 | 0 |
| 3 | A | 14 | 0 | 13 | 0 | 0 |
| 3 | B | 14 | 0 | 13 | 0 | 0 |
| 3 | C | 14 | 0 | 13 | 0 | 0 |
| All | All | 14409 | 0 | 14013 | 425 | 0 |

PRELIMINARY VALIDATION REPORT

The all-atom clashscore is defined as the number of clashes found per 1000 atoms (including hydrogen atoms). The all-atom clashscore for this structure is 15.

All (425) close contacts within the same asymmetric unit are listed below, sorted by their clash magnitude.

| Atom-1 | Atom-2 | Interatomic distance (Å) | Clash overlap (Å) |
|---|---|---|---|
| 1:B:512:VAL:CG1 | 1:C:698:LEU:HD22 | 1.51 | 1.40 |
| 1:B:512:VAL:HG11 | 1:C:698:LEU:CD2 | 1.57 | 1.33 |
| 1:B:439:VAL:HG23 | 1:B:461:LEU:HD11 | 1.27 | 1.16 |
| 1:B:512:VAL:HG21 | 1:C:698:LEU:CD2 | 1.76 | 1.14 |
| 1:A:439:VAL:HG23 | 1:A:461:LEU:HD11 | 1.29 | 1.14 |
| 1:A:513:GLU:OE2 | 1:B:513:GLU:OE2 | 1.65 | 1.13 |
| 1:C:439:VAL:HG23 | 1:C:461:LEU:HD11 | 1.29 | 1.12 |
| 1:C:241:ILE:HB | 2:F:1:NAG:H83 | 1.32 | 1.11 |
| 1:A:441:THR:HB | 1:A:461:LEU:HD23 | 1.28 | 1.10 |
| 1:A:241:ILE:HB | 2:D:1:NAG:H83 | 1.33 | 1.09 |
| 1:B:441:THR:HB | 1:B:461:LEU:HD23 | 1.26 | 1.08 |
| 1:A:513:GLU:OE2 | 1:B:513:GLU:CD | 1.90 | 1.08 |
| 1:B:241:ILE:HB | 2:E:1:NAG:H83 | 1.31 | 1.07 |
| 1:C:441:THR:HB | 1:C:461:LEU:HD23 | 1.28 | 1.05 |
| 1:A:513:GLU:OE1 | 1:B:513:GLU:OE1 | 1.77 | 1.01 |
| 1:A:700:VAL:CG2 | 1:C:512:VAL:HG23 | 1.90 | 1.00 |
| 1:C:512:VAL:O | 1:C:516:MET:HG3 | 1.61 | 1.00 |
| 1:A:513:GLU:CD | 1:B:513:GLU:CD | 2.20 | 0.99 |
| 1:A:700:VAL:HG22 | 1:C:512:VAL:CG2 | 1.95 | 0.97 |
| 1:A:517:LEU:CD1 | 1:B:513:GLU:OE2 | 2.12 | 0.97 |
| 1:B:512:VAL:HG21 | 1:C:698:LEU:HD23 | 1.44 | 0.97 |
| 1:B:678:MET:HE1 | 1:C:134:GLU:HA | 1.48 | 0.95 |
| 1:A:134:GLU:HA | 1:C:678:MET:HE1 | 1.46 | 0.94 |
| 1:B:439:VAL:HG23 | 1:B:461:LEU:CD1 | 1.99 | 0.91 |
| 1:A:439:VAL:HG23 | 1:A:461:LEU:CD1 | 2.02 | 0.90 |
| 1:C:439:VAL:HG23 | 1:C:461:LEU:CD1 | 2.02 | 0.88 |
| 1:A:700:VAL:HG23 | 1:C:512:VAL:HG23 | 1.55 | 0.86 |
| 1:B:512:VAL:CG2 | 1:C:698:LEU:CD2 | 2.52 | 0.86 |
| 1:A:517:LEU:HD11 | 1:B:513:GLU:OE2 | 1.77 | 0.84 |
| 1:B:512:VAL:HG21 | 1:C:698:LEU:HD21 | 1.59 | 0.82 |
| 1:A:626:GLU:OE1 | 1:A:639:ARG:NH2 | 2.13 | 0.82 |
| 1:C:626:GLU:OE1 | 1:C:639:ARG:NH2 | 2.13 | 0.82 |
| 1:A:513:GLU:OE1 | 1:B:513:GLU:CD | 2.18 | 0.82 |
| 1:B:512:VAL:HG23 | 1:C:700:VAL:CG2 | 2.09 | 0.81 |
| 1:B:626:GLU:OE1 | 1:B:639:ARG:NH2 | 2.13 | 0.80 |
| 1:C:241:ILE:HB | 2:F:1:NAG:C8 | 2.10 | 0.80 |
| 1:B:607:ARG:HD2 | 1:B:642:LEU:HB3 | 1.65 | 0.79 |

*Continued on next page...*

PRELIMINARY VALIDATION REPORT

*Continued from previous page...*

| Atom-1 | Atom-2 | Interatomic distance (Å) | Clash overlap (Å) |
|---|---|---|---|
| 1:C:607:ARG:HD2 | 1:C:642:LEU:HB3 | 1.65 | 0.79 |
| 1:A:285:ARG:NH2 | 1:B:710:GLY:O | 2.16 | 0.79 |
| 1:A:241:ILE:HB | 2:D:1:NAG:C8 | 2.11 | 0.78 |
| 1:A:710:GLY:O | 1:C:285:ARG:NH2 | 2.17 | 0.78 |
| 1:B:285:ARG:NH2 | 1:C:710:GLY:O | 2.17 | 0.78 |
| 1:A:374:TRP:CH2 | 1:B:698:LEU:HD21 | 2.18 | 0.78 |
| 1:A:607:ARG:HD2 | 1:A:642:LEU:HB3 | 1.65 | 0.78 |
| 1:B:512:VAL:HG23 | 1:C:700:VAL:HG22 | 1.66 | 0.78 |
| 1:A:678:MET:HE1 | 1:B:134:GLU:HA | 1.64 | 0.77 |
| 1:B:241:ILE:HB | 2:E:1:NAG:C8 | 2.11 | 0.77 |
| 1:B:374:TRP:CH2 | 1:C:698:LEU:HD21 | 2.19 | 0.77 |
| 1:B:374:TRP:CZ2 | 1:C:698:LEU:HD21 | 2.20 | 0.77 |
| 1:A:374:TRP:CZ2 | 1:B:698:LEU:HD21 | 2.21 | 0.76 |
| 1:B:678:MET:HE3 | 1:C:132:ARG:HG3 | 1.69 | 0.74 |
| 1:A:132:ARG:HG3 | 1:C:678:MET:HE3 | 1.70 | 0.73 |
| 1:C:365:ARG:O | 1:C:416:GLN:NE2 | 2.21 | 0.73 |
| 1:A:512:VAL:HG23 | 1:B:700:VAL:HG22 | 1.69 | 0.73 |
| 1:A:365:ARG:O | 1:A:416:GLN:NE2 | 2.21 | 0.73 |
| 1:A:513:GLU:HG3 | 1:C:513:GLU:OE1 | 1.88 | 0.72 |
| 1:A:439:VAL:CG2 | 1:A:461:LEU:HD11 | 2.14 | 0.72 |
| 1:A:534:ARG:NH1 | 1:B:686:ASN:O | 2.21 | 0.72 |
| 1:A:686:ASN:O | 1:C:534:ARG:NH1 | 2.22 | 0.72 |
| 1:B:439:VAL:CG2 | 1:B:461:LEU:HD11 | 2.14 | 0.72 |
| 1:B:365:ARG:O | 1:B:416:GLN:NE2 | 2.21 | 0.71 |
| 1:A:203:ILE:HD13 | 1:A:207:ILE:HD12 | 1.72 | 0.71 |
| 1:C:439:VAL:CG2 | 1:C:461:LEU:HD11 | 2.14 | 0.71 |
| 1:B:534:ARG:NH1 | 1:C:686:ASN:O | 2.23 | 0.70 |
| 1:A:700:VAL:HG22 | 1:C:512:VAL:HG22 | 1.73 | 0.70 |
| 1:A:592:ARG:HE | 1:A:594:ILE:HD11 | 1.57 | 0.69 |
| 1:B:592:ARG:HE | 1:B:594:ILE:HD11 | 1.57 | 0.69 |
| 1:B:203:ILE:HD13 | 1:B:207:ILE:HD12 | 1.72 | 0.69 |
| 1:A:134:GLU:HA | 1:C:678:MET:CE | 2.21 | 0.69 |
| 1:C:203:ILE:HD13 | 1:C:207:ILE:HD12 | 1.72 | 0.69 |
| 1:A:678:MET:CE | 1:B:134:GLU:HA | 2.23 | 0.69 |
| 1:A:698:LEU:HD21 | 1:C:374:TRP:CH2 | 2.28 | 0.69 |
| 1:A:700:VAL:CG2 | 1:C:512:VAL:CG2 | 2.57 | 0.69 |
| 1:C:592:ARG:HE | 1:C:594:ILE:HD11 | 1.57 | 0.68 |
| 1:B:441:THR:HB | 1:B:461:LEU:CD2 | 2.15 | 0.67 |
| 1:A:441:THR:HB | 1:A:461:LEU:CD2 | 2.18 | 0.67 |
| 1:A:679:ILE:HG21 | 1:B:131:VAL:HG21 | 1.76 | 0.66 |
| 1:B:679:ILE:HG21 | 1:C:131:VAL:HG21 | 1.77 | 0.66 |

*Continued on next page...*

PRELIMINARY VALIDATION REPORT

*Continued from previous page...*

| Atom-1 | Atom-2 | Interatomic distance (Å) | Clash overlap (Å) |
|---|---|---|---|
| 1:B:678:MET:CE | 1:C:134:GLU:HA | 2.25 | 0.66 |
| 1:B:439:VAL:O | 1:B:461:LEU:HG | 1.96 | 0.65 |
| 1:A:131:VAL:HG21 | 1:C:679:ILE:HG21 | 1.77 | 0.65 |
| 1:A:439:VAL:O | 1:A:461:LEU:HG | 1.97 | 0.65 |
| 1:C:439:VAL:O | 1:C:461:LEU:HG | 1.96 | 0.65 |
| 1:B:376:GLU:N | 1:B:376:GLU:OE1 | 2.29 | 0.64 |
| 1:A:678:MET:HE3 | 1:B:132:ARG:HG3 | 1.79 | 0.64 |
| 1:A:376:GLU:N | 1:A:376:GLU:OE1 | 2.29 | 0.64 |
| 1:A:698:LEU:HD21 | 1:C:374:TRP:CZ2 | 2.32 | 0.64 |
| 1:A:716:GLU:OE2 | 1:A:719:ARG:NH2 | 2.31 | 0.64 |
| 1:B:512:VAL:CG2 | 1:C:700:VAL:HG22 | 2.28 | 0.64 |
| 1:C:441:THR:HB | 1:C:461:LEU:CD2 | 2.18 | 0.63 |
| 1:B:512:VAL:O | 1:B:516:MET:HG3 | 1.99 | 0.63 |
| 2:E:2:NAG:H61 | 2:E:3:MAN:H2 | 1.79 | 0.63 |
| 2:F:2:NAG:H61 | 2:F:3:MAN:H2 | 1.80 | 0.63 |
| 1:C:660:VAL:HG12 | 1:C:670:GLU:HG3 | 1.81 | 0.63 |
| 1:A:517:LEU:HD12 | 1:B:513:GLU:OE2 | 1.96 | 0.63 |
| 1:C:376:GLU:N | 1:C:376:GLU:OE1 | 2.29 | 0.63 |
| 1:B:716:GLU:OE2 | 1:B:719:ARG:NH2 | 2.31 | 0.63 |
| 1:B:660:VAL:HG12 | 1:B:670:GLU:HG3 | 1.81 | 0.62 |
| 1:C:716:GLU:OE2 | 1:C:719:ARG:NH2 | 2.31 | 0.62 |
| 1:A:680:SER:HA | 1:B:134:GLU:OE1 | 1.99 | 0.62 |
| 1:B:512:VAL:CB | 1:C:698:LEU:CD2 | 2.77 | 0.62 |
| 1:C:512:VAL:O | 1:C:512:VAL:HG12 | 2.00 | 0.62 |
| 1:A:134:GLU:OE1 | 1:C:680:SER:HA | 1.99 | 0.62 |
| 1:A:660:VAL:HG12 | 1:A:670:GLU:HG3 | 1.81 | 0.62 |
| 1:C:364:LYS:HE2 | 1:C:414:LEU:HD21 | 1.81 | 0.62 |
| 1:B:512:VAL:CG2 | 1:C:698:LEU:HD23 | 2.23 | 0.61 |
| 2:D:2:NAG:H61 | 2:D:3:MAN:H2 | 1.81 | 0.61 |
| 1:B:241:ILE:CB | 2:E:1:NAG:H83 | 2.19 | 0.61 |
| 1:B:364:LYS:HE2 | 1:B:414:LEU:HD21 | 1.81 | 0.61 |
| 1:A:364:LYS:HE2 | 1:A:414:LEU:HD21 | 1.81 | 0.61 |
| 1:A:512:VAL:HG23 | 1:B:700:VAL:CG2 | 2.30 | 0.61 |
| 1:C:155:VAL:HG12 | 1:C:455:VAL:HG22 | 1.83 | 0.61 |
| 1:C:241:ILE:CB | 2:F:1:NAG:H83 | 2.20 | 0.61 |
| 1:A:512:VAL:HG12 | 1:A:515:ALA:HB3 | 1.82 | 0.60 |
| 1:A:679:ILE:HG23 | 1:B:131:VAL:HB | 1.85 | 0.59 |
| 1:B:512:VAL:CG1 | 1:C:698:LEU:CD2 | 2.39 | 0.59 |
| 1:A:155:VAL:HG12 | 1:A:455:VAL:HG22 | 1.83 | 0.59 |
| 1:A:534:ARG:HH12 | 1:B:688:THR:HG23 | 1.67 | 0.59 |
| 1:A:131:VAL:HB | 1:C:679:ILE:HG23 | 1.85 | 0.59 |

*Continued on next page...*

PRELIMINARY VALIDATION REPORT

*Continued from previous page...*

| Atom-1 | Atom-2 | Interatomic distance (Å) | Clash overlap (Å) |
|---|---|---|---|
| 1:B:569:ARG:HH22 | 1:B:632:ASP:HB3 | 1.67 | 0.59 |
| 1:B:155:VAL:HG12 | 1:B:455:VAL:HG22 | 1.83 | 0.59 |
| 1:A:678:MET:HE2 | 1:B:134:GLU:N | 2.18 | 0.58 |
| 1:A:513:GLU:CD | 1:B:513:GLU:CG | 2.72 | 0.58 |
| 1:A:569:ARG:HH22 | 1:A:632:ASP:HB3 | 1.67 | 0.58 |
| 1:C:569:ARG:HH22 | 1:C:632:ASP:HB3 | 1.67 | 0.58 |
| 1:C:430:TYR:OH | 1:C:438:HIS:O | 2.22 | 0.58 |
| 1:B:678:MET:HE2 | 1:C:134:GLU:N | 2.19 | 0.57 |
| 1:A:241:ILE:CB | 2:D:1:NAG:H83 | 2.21 | 0.57 |
| 1:A:189:THR:HG21 | 1:A:269:TYR:HE1 | 1.70 | 0.57 |
| 1:B:158:GLU:H | 1:B:508:THR:HG21 | 1.70 | 0.57 |
| 1:A:224:HIS:CE1 | 1:C:735:VAL:HG11 | 2.39 | 0.57 |
| 1:A:430:TYR:OH | 1:A:438:HIS:O | 2.22 | 0.57 |
| 1:B:189:THR:HG21 | 1:B:269:TYR:HE1 | 1.70 | 0.57 |
| 1:A:134:GLU:N | 1:C:678:MET:HE2 | 2.18 | 0.57 |
| 1:A:613:LEU:HD22 | 1:A:639:ARG:HD3 | 1.87 | 0.56 |
| 1:C:158:GLU:H | 1:C:508:THR:HG21 | 1.70 | 0.56 |
| 1:B:511:SER:O | 1:C:700:VAL:HG21 | 2.05 | 0.56 |
| 1:A:158:GLU:H | 1:A:508:THR:HG21 | 1.70 | 0.56 |
| 1:B:680:SER:HA | 1:C:134:GLU:OE1 | 2.04 | 0.56 |
| 1:C:189:THR:HG21 | 1:C:269:TYR:HE1 | 1.70 | 0.56 |
| 1:A:374:TRP:HH2 | 1:B:698:LEU:HD21 | 1.68 | 0.56 |
| 1:B:678:MET:CE | 1:C:134:GLU:N | 2.69 | 0.56 |
| 1:B:430:TYR:OH | 1:B:438:HIS:O | 2.22 | 0.56 |
| 1:C:613:LEU:HD22 | 1:C:639:ARG:HD3 | 1.87 | 0.56 |
| 1:B:613:LEU:HD22 | 1:B:639:ARG:HD3 | 1.87 | 0.56 |
| 1:A:134:GLU:CA | 1:C:678:MET:CE | 2.83 | 0.56 |
| 1:C:335:ASP:HB2 | 1:C:342:LEU:HD21 | 1.88 | 0.55 |
| 1:A:512:VAL:O | 1:A:515:ALA:N | 2.39 | 0.55 |
| 1:A:569:ARG:NH2 | 1:A:632:ASP:HB3 | 2.22 | 0.55 |
| 1:B:534:ARG:HH12 | 1:C:688:THR:HG23 | 1.71 | 0.55 |
| 1:B:374:TRP:HH2 | 1:C:698:LEU:HD21 | 1.70 | 0.55 |
| 1:C:569:ARG:NH2 | 1:C:632:ASP:HB3 | 2.22 | 0.55 |
| 1:A:679:ILE:CG1 | 1:B:132:ARG:O | 2.55 | 0.55 |
| 1:B:679:ILE:HG23 | 1:C:131:VAL:HB | 1.88 | 0.55 |
| 1:C:231:GLU:HG3 | 1:C:260:TYR:CG | 2.42 | 0.55 |
| 1:A:134:GLU:N | 1:C:678:MET:CE | 2.69 | 0.55 |
| 1:A:231:GLU:HG3 | 1:A:260:TYR:CG | 2.42 | 0.55 |
| 1:B:231:GLU:HG3 | 1:B:260:TYR:CG | 2.42 | 0.55 |
| 1:C:589:SER:O | 1:C:591:THR:HG23 | 2.07 | 0.55 |
| 2:D:2:NAG:H5 | 2:D:2:NAG:HN2 | 1.72 | 0.55 |

*Continued on next page...*

PRELIMINARY VALIDATION REPORT

*Continued from previous page...*

| Atom-1 | Atom-2 | Interatomic distance (Å) | Clash overlap (Å) |
|---|---|---|---|
| 1:A:118:THR:OG1 | 1:A:586:GLU:OE1 | 2.21 | 0.54 |
| 1:A:589:SER:O | 1:A:591:THR:HG23 | 2.07 | 0.54 |
| 1:A:678:MET:CE | 1:B:134:GLU:CA | 2.85 | 0.54 |
| 1:B:452:GLY:O | 1:B:508:THR:HG22 | 2.07 | 0.54 |
| 1:B:678:MET:CE | 1:C:134:GLU:CA | 2.85 | 0.54 |
| 1:A:335:ASP:HB2 | 1:A:342:LEU:HD21 | 1.88 | 0.54 |
| 1:A:517:LEU:HD12 | 1:B:513:GLU:OE1 | 2.07 | 0.54 |
| 1:B:569:ARG:NH2 | 1:B:632:ASP:HB3 | 2.22 | 0.54 |
| 1:A:452:GLY:O | 1:A:508:THR:HG22 | 2.07 | 0.54 |
| 1:B:589:SER:O | 1:B:591:THR:HG23 | 2.07 | 0.54 |
| 1:A:688:THR:HG23 | 1:C:534:ARG:HH12 | 1.72 | 0.54 |
| 1:A:132:ARG:O | 1:C:679:ILE:CG1 | 2.55 | 0.54 |
| 1:C:512:VAL:O | 1:C:516:MET:CG | 2.46 | 0.54 |
| 1:A:150:GLU:HG3 | 1:A:151:GLY:N | 2.23 | 0.54 |
| 1:B:735:VAL:HG11 | 1:C:224:HIS:CE1 | 2.43 | 0.54 |
| 1:B:335:ASP:HB2 | 1:B:342:LEU:HD21 | 1.88 | 0.54 |
| 1:A:517:LEU:HD12 | 1:B:513:GLU:CD | 2.27 | 0.54 |
| 1:C:461:LEU:HD12 | 1:C:461:LEU:O | 2.08 | 0.54 |
| 1:B:150:GLU:HG3 | 1:B:151:GLY:N | 2.23 | 0.54 |
| 1:B:118:THR:OG1 | 1:B:586:GLU:OE1 | 2.21 | 0.54 |
| 1:C:150:GLU:HG3 | 1:C:151:GLY:N | 2.23 | 0.54 |
| 1:C:452:GLY:O | 1:C:508:THR:HG22 | 2.07 | 0.53 |
| 1:C:200:VAL:O | 1:C:204:THR:HG22 | 2.09 | 0.53 |
| 1:B:381:VAL:HG23 | 1:B:392:THR:HB | 1.91 | 0.53 |
| 1:A:381:VAL:HG23 | 1:A:392:THR:HB | 1.91 | 0.53 |
| 1:B:461:LEU:HD12 | 1:B:461:LEU:O | 2.08 | 0.53 |
| 1:A:512:VAL:O | 1:A:515:ALA:HB3 | 2.08 | 0.53 |
| 1:B:220:VAL:HG22 | 1:B:225:LYS:HD2 | 1.90 | 0.53 |
| 1:A:678:MET:CE | 1:B:134:GLU:N | 2.72 | 0.53 |
| 1:C:170:TYR:HB3 | 1:C:198:ILE:HD12 | 1.91 | 0.53 |
| 1:C:286:SER:HB2 | 1:C:293:PHE:HB3 | 1.91 | 0.53 |
| 1:C:118:THR:OG1 | 1:C:586:GLU:OE1 | 2.21 | 0.53 |
| 1:A:461:LEU:HD12 | 1:A:461:LEU:O | 2.08 | 0.53 |
| 1:A:220:VAL:HG22 | 1:A:225:LYS:HD2 | 1.91 | 0.52 |
| 1:A:679:ILE:HG12 | 1:B:132:ARG:O | 2.09 | 0.52 |
| 1:B:286:SER:HB2 | 1:B:293:PHE:HB3 | 1.91 | 0.52 |
| 1:A:200:VAL:O | 1:A:204:THR:HG22 | 2.09 | 0.52 |
| 1:B:170:TYR:HB3 | 1:B:198:ILE:HD12 | 1.91 | 0.52 |
| 1:A:286:SER:HB2 | 1:A:293:PHE:HB3 | 1.91 | 0.52 |
| 1:B:601:VAL:HG23 | 1:B:608:CYS:HA | 1.92 | 0.52 |
| 1:A:700:VAL:HG21 | 1:C:511:SER:O | 2.09 | 0.52 |

*Continued on next page...*

PRELIMINARY VALIDATION REPORT

Continued from previous page...

| Atom-1 | Atom-2 | Interatomic distance (Å) | Clash overlap (Å) |
|---|---|---|---|
| 1:A:735:VAL:HG11 | 1:B:224:HIS:CE1 | 2.45 | 0.52 |
| 1:C:220:VAL:HG22 | 1:C:225:LYS:HD2 | 1.91 | 0.52 |
| 1:A:132:ARG:O | 1:C:679:ILE:HG12 | 2.09 | 0.52 |
| 1:A:170:TYR:HB3 | 1:A:198:ILE:HD12 | 1.91 | 0.52 |
| 1:B:590:ASP:OD2 | 1:B:619:LEU:HB2 | 2.10 | 0.52 |
| 1:C:381:VAL:HG23 | 1:C:392:THR:HB | 1.91 | 0.52 |
| 1:C:590:ASP:OD2 | 1:C:619:LEU:HB2 | 2.10 | 0.52 |
| 1:B:300:ILE:HD11 | 1:C:712:LEU:HD11 | 1.92 | 0.51 |
| 1:B:200:VAL:O | 1:B:204:THR:HG22 | 2.09 | 0.51 |
| 1:A:601:VAL:HG23 | 1:A:608:CYS:HA | 1.91 | 0.51 |
| 1:A:590:ASP:OD2 | 1:A:619:LEU:HB2 | 2.10 | 0.51 |
| 1:C:601:VAL:HG23 | 1:C:608:CYS:HA | 1.92 | 0.51 |
| 1:A:230:ASN:OD1 | 1:A:274:SER:HA | 2.11 | 0.51 |
| 1:A:731:ILE:O | 1:B:324:ARG:NH2 | 2.44 | 0.51 |
| 1:C:196:VAL:HG21 | 1:C:219:TYR:CD1 | 2.46 | 0.51 |
| 1:A:196:VAL:HG21 | 1:A:219:TYR:CD1 | 2.46 | 0.51 |
| 1:A:129:THR:OG1 | 1:A:581:VAL:O | 2.29 | 0.50 |
| 1:A:658:HIS:HB3 | 1:A:670:GLU:HG2 | 1.93 | 0.50 |
| 1:A:300:ILE:HD11 | 1:B:712:LEU:HD11 | 1.93 | 0.50 |
| 1:B:398:THR:HG23 | 1:B:515:ALA:HB1 | 1.93 | 0.50 |
| 1:C:230:ASN:OD1 | 1:C:274:SER:HA | 2.11 | 0.50 |
| 1:A:698:LEU:HD21 | 1:C:374:TRP:HH2 | 1.76 | 0.50 |
| 1:B:125:PRO:HD2 | 1:B:573:ARG:NH1 | 2.27 | 0.50 |
| 1:B:196:VAL:HG21 | 1:B:219:TYR:CD1 | 2.46 | 0.50 |
| 1:B:230:ASN:OD1 | 1:B:274:SER:HA | 2.11 | 0.50 |
| 1:C:129:THR:OG1 | 1:C:581:VAL:O | 2.29 | 0.50 |
| 1:B:157:LYS:HG2 | 1:B:374:TRP:HB2 | 1.93 | 0.50 |
| 1:B:702:THR:HG22 | 1:B:703:ARG:H | 1.77 | 0.50 |
| 1:C:125:PRO:HD2 | 1:C:573:ARG:NH1 | 2.27 | 0.50 |
| 1:C:658:HIS:HB3 | 1:C:670:GLU:HG2 | 1.93 | 0.50 |
| 1:A:125:PRO:HD2 | 1:A:573:ARG:NH1 | 2.27 | 0.50 |
| 1:A:678:MET:HE2 | 1:B:134:GLU:CA | 2.41 | 0.50 |
| 1:C:157:LYS:HG2 | 1:C:374:TRP:HB2 | 1.93 | 0.50 |
| 1:A:134:GLU:CA | 1:C:678:MET:HE1 | 2.29 | 0.50 |
| 1:B:129:THR:OG1 | 1:B:581:VAL:O | 2.29 | 0.50 |
| 1:C:383:ASP:OD1 | 1:C:390:ARG:HB3 | 2.12 | 0.50 |
| 1:A:383:ASP:OD1 | 1:A:390:ARG:HB3 | 2.12 | 0.50 |
| 1:A:702:THR:HG22 | 1:A:703:ARG:H | 1.77 | 0.50 |
| 1:C:702:THR:HG22 | 1:C:703:ARG:H | 1.76 | 0.50 |
| 1:B:650:LYS:HG2 | 1:B:663:GLU:OE2 | 2.12 | 0.50 |
| 1:B:658:HIS:HB3 | 1:B:670:GLU:HG2 | 1.93 | 0.50 |

Continued on next page...

PRELIMINARY VALIDATION REPORT

*Continued from previous page...*

| Atom-1 | Atom-2 | Interatomic distance (Å) | Clash overlap (Å) |
|---|---|---|---|
| 1:C:265:THR:O | 1:C:268:THR:HG22 | 2.12 | 0.50 |
| 1:B:383:ASP:OD1 | 1:B:390:ARG:HB3 | 2.12 | 0.49 |
| 1:A:265:THR:O | 1:A:268:THR:HG22 | 2.12 | 0.49 |
| 1:B:265:THR:O | 1:B:268:THR:HG22 | 2.12 | 0.49 |
| 2:F:2:NAG:H5 | 2:F:2:NAG:HN2 | 1.76 | 0.49 |
| 1:A:157:LYS:HG2 | 1:A:374:TRP:HB2 | 1.93 | 0.49 |
| 1:A:650:LYS:HG2 | 1:A:663:GLU:OE2 | 2.12 | 0.49 |
| 1:A:712:LEU:HD11 | 1:C:300:ILE:HD11 | 1.93 | 0.49 |
| 1:A:244:LYS:HA | 2:D:2:NAG:O3 | 2.13 | 0.49 |
| 1:C:650:LYS:HG2 | 1:C:663:GLU:OE2 | 2.12 | 0.49 |
| 1:B:125:PRO:HB3 | 1:B:582:SER:HB3 | 1.95 | 0.48 |
| 1:B:441:THR:CB | 1:B:461:LEU:HD23 | 2.20 | 0.48 |
| 1:A:131:VAL:CB | 1:C:679:ILE:HG23 | 2.44 | 0.48 |
| 1:B:679:ILE:CG1 | 1:C:132:ARG:O | 2.61 | 0.48 |
| 1:C:125:PRO:HB3 | 1:C:582:SER:HB3 | 1.95 | 0.48 |
| 1:B:374:TRP:HZ2 | 1:C:698:LEU:HD21 | 1.76 | 0.48 |
| 1:A:391:PHE:HE1 | 1:A:402:SER:OG | 1.97 | 0.48 |
| 1:C:733:LYS:HE3 | 1:C:733:LYS:HB2 | 1.66 | 0.48 |
| 1:A:324:ARG:NH2 | 1:C:731:ILE:O | 2.47 | 0.47 |
| 1:B:685:LEU:HG | 1:B:687:LEU:HG | 1.96 | 0.47 |
| 1:C:391:PHE:HE1 | 1:C:402:SER:OG | 1.97 | 0.47 |
| 1:C:578:VAL:HG12 | 1:C:579:ILE:N | 2.29 | 0.47 |
| 1:C:513:GLU:C | 1:C:515:ALA:N | 2.67 | 0.47 |
| 1:A:164:LYS:NZ | 1:A:285:ARG:HH11 | 2.12 | 0.47 |
| 1:A:125:PRO:HB3 | 1:A:582:SER:HB3 | 1.95 | 0.47 |
| 1:C:172:LYS:O | 1:C:196:VAL:HG12 | 2.15 | 0.47 |
| 1:A:613:LEU:HD23 | 1:A:628:GLN:HB3 | 1.96 | 0.47 |
| 1:B:164:LYS:NZ | 1:B:285:ARG:HH11 | 2.12 | 0.47 |
| 1:A:172:LYS:O | 1:A:196:VAL:HG12 | 2.15 | 0.47 |
| 1:B:164:LYS:HG2 | 1:B:285:ARG:HG2 | 1.97 | 0.47 |
| 1:B:437:SER:OG | 1:B:438:HIS:ND1 | 2.32 | 0.47 |
| 1:A:733:LYS:HE3 | 1:A:733:LYS:HB2 | 1.66 | 0.47 |
| 1:B:391:PHE:HE1 | 1:B:402:SER:OG | 1.97 | 0.47 |
| 1:B:578:VAL:HG12 | 1:B:579:ILE:N | 2.29 | 0.47 |
| 1:B:172:LYS:O | 1:B:196:VAL:HG12 | 2.15 | 0.47 |
| 1:A:578:VAL:HG12 | 1:A:579:ILE:N | 2.29 | 0.47 |
| 2:E:2:NAG:HN2 | 2:E:2:NAG:H5 | 1.79 | 0.47 |
| 1:A:290:TYR:CZ | 1:A:368:VAL:HG21 | 2.51 | 0.46 |
| 1:B:512:VAL:HG23 | 1:C:700:VAL:HG23 | 1.94 | 0.46 |
| 1:A:685:LEU:HG | 1:A:687:LEU:HG | 1.96 | 0.46 |
| 1:C:613:LEU:HD23 | 1:C:628:GLN:HB3 | 1.96 | 0.46 |

*Continued on next page...*

PRELIMINARY VALIDATION REPORT

Continued from previous page...

| Atom-1 | Atom-2 | Interatomic distance (Å) | Clash overlap (Å) |
|---|---|---|---|
| 1:B:731:ILE:O | 1:C:324:ARG:NH2 | 2.49 | 0.46 |
| 1:C:164:LYS:HG2 | 1:C:285:ARG:HG2 | 1.97 | 0.46 |
| 1:C:164:LYS:NZ | 1:C:285:ARG:HH11 | 2.12 | 0.46 |
| 1:A:186:THR:O | 1:C:226:VAL:HG13 | 2.15 | 0.46 |
| 1:A:679:ILE:HG23 | 1:B:131:VAL:CB | 2.44 | 0.46 |
| 1:B:613:LEU:HD23 | 1:B:628:GLN:HB3 | 1.96 | 0.46 |
| 1:A:164:LYS:HG2 | 1:A:285:ARG:HG2 | 1.97 | 0.46 |
| 1:A:627:GLY:O | 1:A:639:ARG:NE | 2.48 | 0.46 |
| 1:A:517:LEU:CG | 1:B:513:GLU:OE2 | 2.61 | 0.46 |
| 1:A:517:LEU:CD1 | 1:B:513:GLU:CD | 2.79 | 0.46 |
| 1:B:290:TYR:CZ | 1:B:368:VAL:HG21 | 2.51 | 0.46 |
| 1:B:679:ILE:HG12 | 1:C:132:ARG:O | 2.15 | 0.46 |
| 1:B:512:VAL:HG11 | 1:C:698:LEU:HD22 | 0.62 | 0.46 |
| 1:A:226:VAL:HG13 | 1:B:186:THR:O | 2.16 | 0.46 |
| 1:A:679:ILE:HG13 | 1:B:132:ARG:O | 2.15 | 0.46 |
| 1:A:735:VAL:HG12 | 1:B:223:ASN:O | 2.16 | 0.46 |
| 1:C:290:TYR:CZ | 1:C:368:VAL:HG21 | 2.51 | 0.46 |
| 1:B:158:GLU:HG2 | 1:B:159:ASN:N | 2.31 | 0.46 |
| 1:C:158:GLU:HG2 | 1:C:159:ASN:N | 2.31 | 0.46 |
| 1:A:513:GLU:CD | 1:B:513:GLU:HG2 | 2.36 | 0.45 |
| 1:B:679:ILE:HG23 | 1:C:131:VAL:CB | 2.45 | 0.45 |
| 1:C:685:LEU:HG | 1:C:687:LEU:HG | 1.96 | 0.45 |
| 1:B:448:LEU:HD12 | 1:B:453:PHE:O | 2.16 | 0.45 |
| 1:C:448:LEU:HD12 | 1:C:453:PHE:O | 2.16 | 0.45 |
| 1:A:132:ARG:O | 1:C:679:ILE:HG13 | 2.17 | 0.45 |
| 1:B:301:ILE:HD12 | 1:B:352:THR:HG21 | 1.98 | 0.45 |
| 1:B:627:GLY:O | 1:B:639:ARG:NE | 2.48 | 0.45 |
| 1:C:592:ARG:O | 1:C:617:VAL:HG22 | 2.17 | 0.45 |
| 1:B:439:VAL:CG2 | 1:B:461:LEU:CD1 | 2.84 | 0.45 |
| 1:A:448:LEU:HD12 | 1:A:453:PHE:O | 2.16 | 0.45 |
| 1:B:441:THR:HG22 | 1:B:459:PRO:O | 2.17 | 0.45 |
| 1:A:374:TRP:HZ2 | 1:B:698:LEU:HD21 | 1.78 | 0.45 |
| 1:B:652:TYR:N | 1:C:575:LEU:O | 2.37 | 0.45 |
| 1:A:592:ARG:O | 1:A:617:VAL:HG22 | 2.17 | 0.45 |
| 1:B:226:VAL:HG13 | 1:C:186:THR:O | 2.17 | 0.45 |
| 1:B:678:MET:CE | 1:C:133:LEU:C | 2.85 | 0.45 |
| 1:C:513:GLU:C | 1:C:515:ALA:H | 2.21 | 0.45 |
| 1:A:341:LEU:HA | 1:A:341:LEU:HD23 | 1.80 | 0.45 |
| 1:A:301:ILE:HD12 | 1:A:352:THR:HG21 | 1.98 | 0.45 |
| 1:A:513:GLU:OE2 | 1:B:513:GLU:CG | 2.64 | 0.45 |
| 1:A:441:THR:HG22 | 1:A:459:PRO:O | 2.17 | 0.44 |

Continued on next page...

PRELIMINARY VALIDATION REPORT

*Continued from previous page...*

| Atom-1 | Atom-2 | Interatomic distance (Å) | Clash overlap (Å) |
|---|---|---|---|
| 1:A:691:LYS:HA | 1:A:691:LYS:HD3 | 1.75 | 0.44 |
| 1:A:174:VAL:HG13 | 1:A:194:ASP:HB3 | 1.99 | 0.44 |
| 1:B:592:ARG:O | 1:B:617:VAL:HG22 | 2.17 | 0.44 |
| 1:B:382:ARG:HD2 | 1:B:389:PHE:CD1 | 2.53 | 0.44 |
| 1:C:301:ILE:HD12 | 1:C:352:THR:HG21 | 1.98 | 0.44 |
| 1:C:244:LYS:HA | 2:F:2:NAG:O3 | 2.16 | 0.44 |
| 1:C:441:THR:HG22 | 1:C:459:PRO:O | 2.17 | 0.44 |
| 1:A:382:ARG:HD2 | 1:A:389:PHE:CD1 | 2.53 | 0.44 |
| 1:C:174:VAL:HG13 | 1:C:194:ASP:HB3 | 1.99 | 0.44 |
| 1:C:691:LYS:HD3 | 1:C:691:LYS:HA | 1.75 | 0.44 |
| 1:A:133:LEU:C | 1:C:678:MET:CE | 2.86 | 0.44 |
| 1:A:158:GLU:HG2 | 1:A:159:ASN:N | 2.31 | 0.43 |
| 1:A:661:TYR:HB3 | 1:A:669:ARG:HG2 | 2.00 | 0.43 |
| 1:A:517:LEU:HG | 1:B:513:GLU:OE2 | 2.18 | 0.43 |
| 1:A:227:GLU:HG2 | 1:B:185:TYR:HB2 | 2.00 | 0.43 |
| 1:B:512:VAL:HG12 | 1:B:512:VAL:O | 2.18 | 0.43 |
| 1:C:383:ASP:N | 1:C:383:ASP:OD1 | 2.44 | 0.43 |
| 1:C:627:GLY:O | 1:C:639:ARG:NE | 2.48 | 0.43 |
| 1:C:661:TYR:HB3 | 1:C:669:ARG:HG2 | 2.00 | 0.43 |
| 1:C:341:LEU:HD23 | 1:C:341:LEU:HA | 1.80 | 0.43 |
| 2:D:2:NAG:H5 | 2:D:2:NAG:N2 | 2.34 | 0.43 |
| 1:A:124:PRO:HA | 1:A:125:PRO:HD3 | 1.84 | 0.43 |
| 1:B:735:VAL:HG12 | 1:C:223:ASN:O | 2.19 | 0.43 |
| 1:C:382:ARG:HD2 | 1:C:389:PHE:CD1 | 2.53 | 0.43 |
| 1:B:174:VAL:HG13 | 1:B:194:ASP:HB3 | 1.99 | 0.43 |
| 1:B:227:GLU:HG2 | 1:C:185:TYR:HB2 | 2.01 | 0.43 |
| 1:C:437:SER:HG | 1:C:438:HIS:CE1 | 2.29 | 0.43 |
| 2:E:3:MAN:H62 | 2:E:4:MAN:H2 | 1.27 | 0.43 |
| 1:B:517:LEU:HD23 | 1:B:517:LEU:HA | 1.88 | 0.42 |
| 1:B:661:TYR:HB3 | 1:B:669:ARG:HG2 | 2.00 | 0.42 |
| 1:C:663:GLU:CB | 1:C:668:VAL:HG21 | 2.49 | 0.42 |
| 1:B:663:GLU:CB | 1:B:668:VAL:HG21 | 2.49 | 0.42 |
| 1:A:437:SER:OG | 1:A:438:HIS:ND1 | 2.32 | 0.42 |
| 1:B:611:ARG:HH12 | 1:B:631:THR:HG22 | 1.85 | 0.42 |
| 1:B:196:VAL:CG2 | 1:B:197:PRO:HD2 | 2.50 | 0.42 |
| 1:C:661:TYR:OH | 1:C:663:GLU:OE2 | 2.26 | 0.42 |
| 1:A:223:ASN:O | 1:C:735:VAL:HG12 | 2.18 | 0.42 |
| 1:A:309:LEU:HB2 | 1:A:310:ARG:HH11 | 1.84 | 0.42 |
| 1:A:439:VAL:CG2 | 1:A:461:LEU:CD1 | 2.85 | 0.42 |
| 1:A:512:VAL:CG2 | 1:B:700:VAL:HG22 | 2.44 | 0.42 |
| 1:A:196:VAL:CG2 | 1:A:197:PRO:HD2 | 2.50 | 0.42 |

*Continued on next page...*

PRELIMINARY VALIDATION REPORT

*Continued from previous page...*

| Atom-1 | Atom-2 | Interatomic distance (Å) | Clash overlap (Å) |
|---|---|---|---|
| 1:A:416:GLN:N | 1:A:416:GLN:OE1 | 2.48 | 0.42 |
| 1:A:713:ASP:O | 1:A:717:ILE:HG12 | 2.20 | 0.42 |
| 1:B:254:HIS:HA | 1:B:277:CYS:O | 2.20 | 0.42 |
| 2:F:3:MAN:H62 | 2:F:4:MAN:H2 | 1.38 | 0.42 |
| 1:A:663:GLU:CB | 1:A:668:VAL:HG21 | 2.49 | 0.42 |
| 1:A:679:ILE:CG2 | 1:B:131:VAL:HG21 | 2.48 | 0.42 |
| 1:B:702:THR:HG22 | 1:B:703:ARG:N | 2.35 | 0.42 |
| 1:C:254:HIS:HA | 1:C:277:CYS:O | 2.20 | 0.42 |
| 2:D:3:MAN:H62 | 2:D:4:MAN:H2 | 1.35 | 0.42 |
| 1:A:185:TYR:HB2 | 1:C:227:GLU:HG2 | 2.02 | 0.42 |
| 1:B:380:VAL:HG12 | 1:B:393:MET:HG2 | 2.02 | 0.42 |
| 1:B:679:ILE:CG2 | 1:C:131:VAL:HG21 | 2.46 | 0.42 |
| 1:C:196:VAL:CG2 | 1:C:197:PRO:HD2 | 2.50 | 0.42 |
| 1:C:702:THR:HG22 | 1:C:703:ARG:N | 2.35 | 0.41 |
| 1:C:713:ASP:O | 1:C:717:ILE:HG12 | 2.20 | 0.41 |
| 1:A:546:GLU:HG2 | 1:A:550:TRP:HD1 | 1.85 | 0.41 |
| 1:B:309:LEU:HB2 | 1:B:310:ARG:HH11 | 1.84 | 0.41 |
| 1:B:713:ASP:O | 1:B:717:ILE:HG12 | 2.20 | 0.41 |
| 1:C:437:SER:OG | 1:C:438:HIS:ND1 | 2.32 | 0.41 |
| 1:A:382:ARG:NH1 | 1:A:447:TYR:OH | 2.54 | 0.41 |
| 1:B:578:VAL:CG1 | 1:B:579:ILE:N | 2.84 | 0.41 |
| 1:C:382:ARG:NH1 | 1:C:447:TYR:OH | 2.54 | 0.41 |
| 1:C:578:VAL:CG1 | 1:C:579:ILE:N | 2.84 | 0.41 |
| 1:C:611:ARG:HH12 | 1:C:631:THR:HG22 | 1.85 | 0.41 |
| 1:A:196:VAL:HG22 | 1:A:197:PRO:HD2 | 2.03 | 0.41 |
| 1:A:702:THR:HG22 | 1:A:703:ARG:N | 2.35 | 0.41 |
| 1:A:190:ASN:ND2 | 1:B:190:ASN:OD1 | 2.38 | 0.41 |
| 1:B:139:CYS:HB3 | 1:B:540:CYS:HB2 | 2.00 | 0.41 |
| 1:B:660:VAL:CG1 | 1:B:670:GLU:HG3 | 2.50 | 0.41 |
| 1:B:244:LYS:HA | 2:E:2:NAG:O3 | 2.20 | 0.41 |
| 1:C:439:VAL:CG2 | 1:C:461:LEU:CD1 | 2.85 | 0.41 |
| 1:B:433:ARG:HB3 | 1:B:434:TYR:CD1 | 2.56 | 0.41 |
| 1:B:546:GLU:HG2 | 1:B:550:TRP:HD1 | 1.86 | 0.41 |
| 1:C:433:ARG:HB3 | 1:C:434:TYR:CD1 | 2.56 | 0.41 |
| 1:B:123:PRO:HD2 | 1:B:585:PRO:HD2 | 2.03 | 0.41 |
| 1:C:546:GLU:HG2 | 1:C:550:TRP:HD1 | 1.85 | 0.41 |
| 1:A:123:PRO:HD2 | 1:A:585:PRO:HD2 | 2.03 | 0.41 |
| 1:B:513:GLU:C | 1:B:515:ALA:N | 2.73 | 0.41 |
| 1:B:592:ARG:NE | 1:B:594:ILE:HD11 | 2.31 | 0.41 |
| 1:B:513:GLU:HB3 | 1:C:513:GLU:OE2 | 2.21 | 0.41 |
| 1:B:382:ARG:NH1 | 1:B:447:TYR:OH | 2.54 | 0.41 |

*Continued on next page...*

PRELIMINARY VALIDATION REPORT

*Continued from previous page...*

| Atom-1 | Atom-2 | Interatomic distance (Å) | Clash overlap (Å) |
|---|---|---|---|
| 1:A:254:HIS:HA | 1:A:277:CYS:O | 2.20 | 0.41 |
| 1:A:380:VAL:HG12 | 1:A:393:MET:HG2 | 2.02 | 0.41 |
| 1:C:196:VAL:HG22 | 1:C:197:PRO:HD2 | 2.03 | 0.41 |
| 1:C:309:LEU:HB2 | 1:C:310:ARG:HH11 | 1.84 | 0.41 |
| 1:A:514:PHE:CE1 | 1:C:697:PRO:HB3 | 2.56 | 0.40 |
| 1:A:587:LEU:HD13 | 1:A:635:LEU:HD12 | 2.04 | 0.40 |
| 1:A:678:MET:CE | 1:B:132:ARG:HG3 | 2.50 | 0.40 |
| 1:B:196:VAL:HG22 | 1:B:197:PRO:HD2 | 2.03 | 0.40 |
| 1:A:433:ARG:HB3 | 1:A:434:TYR:CD1 | 2.56 | 0.40 |
| 1:C:229:PHE:HE2 | 1:C:235:PRO:HG3 | 1.87 | 0.40 |
| 1:C:123:PRO:HD2 | 1:C:585:PRO:HD2 | 2.03 | 0.40 |
| 2:F:2:NAG:N2 | 2:F:2:NAG:H5 | 2.36 | 0.40 |
| 1:A:611:ARG:HH12 | 1:A:631:THR:HG22 | 1.85 | 0.40 |
| 1:B:229:PHE:HE2 | 1:B:235:PRO:HG3 | 1.86 | 0.40 |
| 1:C:517:LEU:HD23 | 1:C:517:LEU:HA | 1.88 | 0.40 |
| 1:C:609:TYR:HA | 1:C:642:LEU:HD13 | 2.04 | 0.40 |
| 1:A:578:VAL:CG1 | 1:A:579:ILE:N | 2.84 | 0.40 |
| 1:B:130:ILE:HD12 | 1:B:578:VAL:HG11 | 2.03 | 0.40 |
| 1:C:130:ILE:HD12 | 1:C:578:VAL:HG11 | 2.03 | 0.40 |

There are no symmetry-related clashes.

#### 5.3 Torsion angles (i)

##### 5.3.1 Protein backbone (i)

In the following table, the Percentiles column shows the percent Ramachandran outliers of the chain as a percentile score with respect to all PDB entries followed by that with respect to all EM entries.

The Analysed column shows the number of residues for which the backbone conformation was analysed, and the total number of residues.

| Mol | Chain | Analysed | Favoured | Allowed | Outliers | Percentiles | |
|---|---|---|---|---|---|---|---|
| 1 | A | 580/931 (62%) | 569 (98%) | 11 (2%) | 0 | 100 | 100 |
| 1 | B | 580/931 (62%) | 567 (98%) | 13 (2%) | 0 | 100 | 100 |
| 1 | C | 580/931 (62%) | 567 (98%) | 13 (2%) | 0 | 100 | 100 |
| All | All | 1740/2793 (62%) | 1703 (98%) | 37 (2%) | 0 | 100 | 100 |

There are no Ramachandran outliers to report.

PRELIMINARY VALIDATION REPORT

##### 5.3.2 Protein sidechains (i)

In the following table, the Percentiles column shows the percent sidechain outliers of the chain as a percentile score with respect to all PDB entries followed by that with respect to all EM entries.

The Analysed column shows the number of residues for which the sidechain conformation was analysed, and the total number of residues.

| Mol | Chain | Analysed | Rotameric | Outliers | Percentiles | |
|---|---|---|---|---|---|---|
| 1 | A | 524/815 (64%) | 522 (100%) | 2 (0%) | 91 | 94 |
| 1 | B | 524/815 (64%) | 522 (100%) | 2 (0%) | 91 | 94 |
| 1 | C | 524/815 (64%) | 522 (100%) | 2 (0%) | 91 | 94 |
| All | All | 1572/2445 (64%) | 1566 (100%) | 6 (0%) | 91 | 94 |

All (6) residues with a non-rotameric sidechain are listed below:

| Mol | Chain | Res | Type |
|---|---|---|---|
| 1 | A | 139 | CYS |
| 1 | A | 608 | CYS |
| 1 | B | 139 | CYS |
| 1 | B | 608 | CYS |
| 1 | C | 139 | CYS |
| 1 | C | 608 | CYS |

Some sidechains can be flipped to improve hydrogen bonding and reduce clashes. All (16) such sidechains are listed below:

| Mol | Chain | Res | Type |
|---|---|---|---|
| 1 | A | 223 | ASN |
| 1 | A | 525 | GLN |
| 1 | A | 527 | HIS |
| 1 | A | 543 | GLN |
| 1 | A | 628 | GLN |
| 1 | B | 223 | ASN |
| 1 | B | 525 | GLN |
| 1 | B | 527 | HIS |
| 1 | B | 543 | GLN |
| 1 | B | 628 | GLN |
| 1 | C | 223 | ASN |
| 1 | C | 411 | GLN |
| 1 | C | 525 | GLN |
| 1 | C | 527 | HIS |
| 1 | C | 543 | GLN |

*Continued on next page...*

PRELIMINARY VALIDATION REPORT

*Continued from previous page...*

| Mol | Chain | Res | Type |
|---|---|---|---|
| 1 | C | 628 | GLN |

##### 5.3.3 RNA (i)

There are no RNA molecules in this entry.

#### 5.4 Non-standard residues in protein, DNA, RNA chains (i)

There are no non-standard protein/DNA/RNA residues in this entry.

#### 5.5 Carbohydrates (i)

18 monosaccharides are modelled in this entry.

In the following table, the Counts columns list the number of bonds (or angles) for which Mogul statistics could be retrieved, the number of bonds (or angles) that are observed in the model and the number of bonds (or angles) that are defined in the Chemical Component Dictionary. The Link column lists molecule types, if any, to which the group is linked. The Z score for a bond length (or angle) is the number of standard deviations the observed value is removed from the expected value. A bond length (or angle) with $|Z| > 2$ is considered an outlier worth inspection. RMSZ is the root-mean-square of all Z scores of the bond lengths (or angles).

| Mol | Type | Chain | Res | Link | Bond lengths | | | Bond angles | | |
|---|---|---|---|---|---|---|---|---|---|---|
| | | | | | Counts | RMSZ | $\#\|Z\| > 2$ | Counts | RMSZ | $\#\|Z\| > 2$ |
| 2 | NAG | D | 1 | 1,2 | 14,14,15 | 0.31 | 0 | 17,19,21 | 0.60 | 0 |
| 2 | NAG | D | 2 | 2 | 14,14,15 | 0.29 | 0 | 17,19,21 | 0.71 | 1 (5%) |
| 2 | MAN | D | 3 | 2 | 11,11,12 | 0.23 | 0 | 15,15,17 | 0.70 | 0 |
| 2 | MAN | D | 4 | 2 | 11,11,12 | 0.27 | 0 | 15,15,17 | 0.69 | 0 |
| 2 | MAN | D | 5 | 2 | 11,11,12 | 0.26 | 0 | 15,15,17 | 0.62 | 0 |
| 2 | MAN | D | 6 | 2 | 11,11,12 | 0.26 | 0 | 15,15,17 | 0.63 | 0 |
| 2 | NAG | E | 1 | 1,2 | 14,14,15 | 0.31 | 0 | 17,19,21 | 0.62 | 0 |
| 2 | NAG | E | 2 | 2 | 14,14,15 | 0.29 | 0 | 17,19,21 | 0.73 | 1 (5%) |
| 2 | MAN | E | 3 | 2 | 11,11,12 | 0.22 | 0 | 15,15,17 | 0.68 | 0 |
| 2 | MAN | E | 4 | 2 | 11,11,12 | 0.32 | 0 | 15,15,17 | 0.92 | 1 (6%) |
| 2 | MAN | E | 5 | 2 | 11,11,12 | 0.26 | 0 | 15,15,17 | 0.63 | 0 |
| 2 | MAN | E | 6 | 2 | 11,11,12 | 0.26 | 0 | 15,15,17 | 0.63 | 0 |
| 2 | NAG | F | 1 | 1,2 | 14,14,15 | 0.31 | 0 | 17,19,21 | 0.62 | 0 |
| 2 | NAG | F | 2 | 2 | 14,14,15 | 0.30 | 0 | 17,19,21 | 0.71 | 1 (5%) |
| 2 | MAN | F | 3 | 2 | 11,11,12 | 0.22 | 0 | 15,15,17 | 0.71 | 0 |
| 2 | MAN | F | 4 | 2 | 11,11,12 | 0.27 | 0 | 15,15,17 | 0.72 | 0 |
| 2 | MAN | F | 5 | 2 | 11,11,12 | 0.26 | 0 | 15,15,17 | 0.66 | 0 |

PRELIMINARY VALIDATION REPORT

| Mol | Type | Chain | Res | Link | Bond lengths | | | Bond angles | | |
|---|---|---|---|---|---|---|---|---|---|---|
| | | | | | Counts | RMSZ | #\|Z\| > 2 | Counts | RMSZ | #\|Z\| > 2 |
| 2 | MAN | F | 6 | 2 | 11,11,12 | 0.26 | 0 | 15,15,17 | 0.63 | 0 |

In the following table, the Chirals column lists the number of chiral outliers, the number of chiral centers analysed, the number of these observed in the model and the number defined in the Chemical Component Dictionary. Similar counts are reported in the Torsion and Rings columns. '-' means no outliers of that kind were identified.

| Mol | Type | Chain | Res | Link | Chirals | Torsions | Rings |
|---|---|---|---|---|---|---|---|
| 2 | NAG | D | 1 | 1,2 | - | 0/6/23/26 | 0/1/1/1 |
| 2 | NAG | D | 2 | 2 | - | 2/6/23/26 | 0/1/1/1 |
| 2 | MAN | D | 3 | 2 | - | 1/2/19/22 | 0/1/1/1 |
| 2 | MAN | D | 4 | 2 | - | 2/2/19/22 | 0/1/1/1 |
| 2 | MAN | D | 5 | 2 | - | 1/2/19/22 | 0/1/1/1 |
| 2 | MAN | D | 6 | 2 | - | 2/2/19/22 | 0/1/1/1 |
| 2 | NAG | E | 1 | 1,2 | - | 0/6/23/26 | 0/1/1/1 |
| 2 | NAG | E | 2 | 2 | - | 2/6/23/26 | 0/1/1/1 |
| 2 | MAN | E | 3 | 2 | - | 0/2/19/22 | 0/1/1/1 |
| 2 | MAN | E | 4 | 2 | - | 1/2/19/22 | 0/1/1/1 |
| 2 | MAN | E | 5 | 2 | - | 2/2/19/22 | 0/1/1/1 |
| 2 | MAN | E | 6 | 2 | - | 2/2/19/22 | 0/1/1/1 |
| 2 | NAG | F | 1 | 1,2 | - | 0/6/23/26 | 0/1/1/1 |
| 2 | NAG | F | 2 | 2 | - | 2/6/23/26 | 0/1/1/1 |
| 2 | MAN | F | 3 | 2 | - | 1/2/19/22 | 0/1/1/1 |
| 2 | MAN | F | 4 | 2 | - | 2/2/19/22 | 0/1/1/1 |
| 2 | MAN | F | 5 | 2 | - | 1/2/19/22 | 0/1/1/1 |
| 2 | MAN | F | 6 | 2 | - | 2/2/19/22 | 0/1/1/1 |

There are no bond length outliers.

All (4) bond angle outliers are listed below:

| Mol | Chain | Res | Type | Atoms | Z | Observed($^o$) | Ideal($^o$) |
|---|---|---|---|---|---|---|---|
| 2 | E | 4 | MAN | C1-C2-C3 | 2.84 | 113.16 | 109.67 |
| 2 | E | 2 | NAG | C1-O5-C5 | 2.29 | 115.30 | 112.19 |
| 2 | F | 2 | NAG | C1-O5-C5 | 2.26 | 115.25 | 112.19 |
| 2 | D | 2 | NAG | C1-O5-C5 | 2.22 | 115.20 | 112.19 |

There are no chirality outliers.

All (23) torsion outliers are listed below:

PRELIMINARY VALIDATION REPORT

| Mol | Chain | Res | Type | Atoms |
|---|---|---|---|---|
| 2 | E | 2 | NAG | C8-C7-N2-C2 |
| 2 | E | 2 | NAG | O7-C7-N2-C2 |
| 2 | F | 2 | NAG | C8-C7-N2-C2 |
| 2 | F | 2 | NAG | O7-C7-N2-C2 |
| 2 | E | 6 | MAN | O5-C5-C6-O6 |
| 2 | D | 4 | MAN | O5-C5-C6-O6 |
| 2 | D | 6 | MAN | O5-C5-C6-O6 |
| 2 | F | 4 | MAN | O5-C5-C6-O6 |
| 2 | F | 6 | MAN | O5-C5-C6-O6 |
| 2 | E | 5 | MAN | O5-C5-C6-O6 |
| 2 | D | 6 | MAN | C4-C5-C6-O6 |
| 2 | E | 6 | MAN | C4-C5-C6-O6 |
| 2 | D | 4 | MAN | C4-C5-C6-O6 |
| 2 | F | 4 | MAN | C4-C5-C6-O6 |
| 2 | F | 6 | MAN | C4-C5-C6-O6 |
| 2 | D | 2 | NAG | C8-C7-N2-C2 |
| 2 | F | 5 | MAN | O5-C5-C6-O6 |
| 2 | D | 2 | NAG | O7-C7-N2-C2 |
| 2 | E | 5 | MAN | C4-C5-C6-O6 |
| 2 | D | 5 | MAN | O5-C5-C6-O6 |
| 2 | E | 4 | MAN | C4-C5-C6-O6 |
| 2 | F | 3 | MAN | C4-C5-C6-O6 |
| 2 | D | 3 | MAN | C4-C5-C6-O6 |

There are no ring outliers.

12 monomers are involved in 23 short contacts:

| Mol | Chain | Res | Type | Clashes | Symm-Clashes |
|---|---|---|---|---|---|
| 2 | E | 2 | NAG | 3 | 0 |
| 2 | D | 3 | MAN | 2 | 0 |
| 2 | F | 4 | MAN | 1 | 0 |
| 2 | F | 3 | MAN | 2 | 0 |
| 2 | F | 2 | NAG | 4 | 0 |
| 2 | E | 3 | MAN | 2 | 0 |
| 2 | D | 2 | NAG | 4 | 0 |
| 2 | E | 4 | MAN | 1 | 0 |
| 2 | E | 1 | NAG | 3 | 0 |
| 2 | F | 1 | NAG | 3 | 0 |
| 2 | D | 1 | NAG | 3 | 0 |
| 2 | D | 4 | MAN | 1 | 0 |

The following is a two-dimensional graphical depiction of Mogul quality analysis of bond lengths, bond angles, torsion angles, and ring geometry for oligosaccharide.

PRELIMINARY VALIDATION REPORT

**Oligosaccharide Chain D**

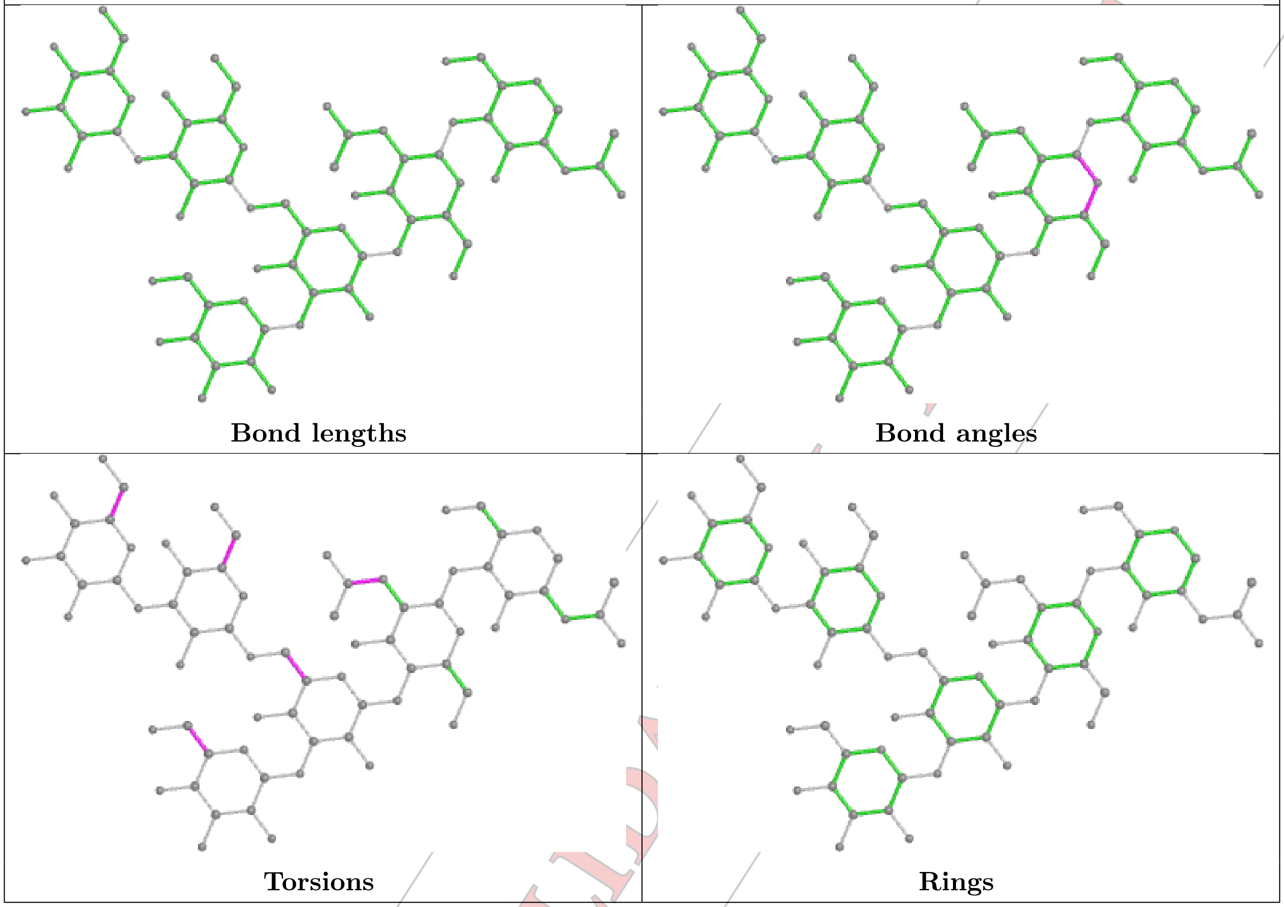

Bond lengths

Bond angles

Torsions

Rings

**Oligosaccharide Chain E**

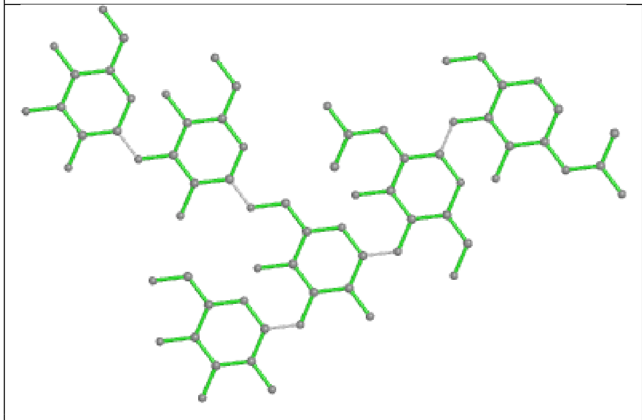

**Bond lengths**

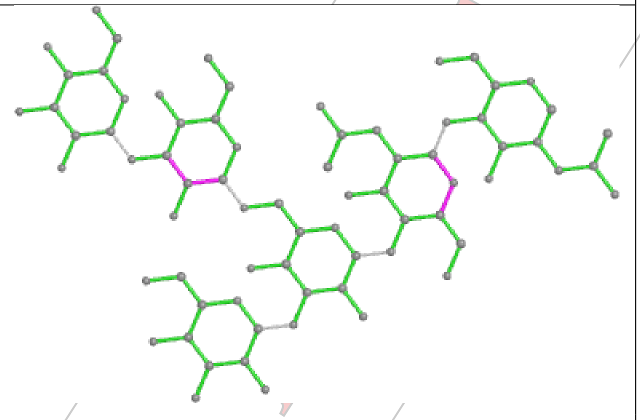

**Bond angles**

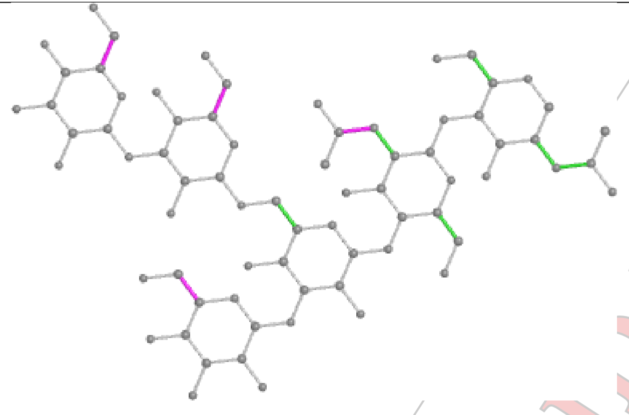

**Torsions**

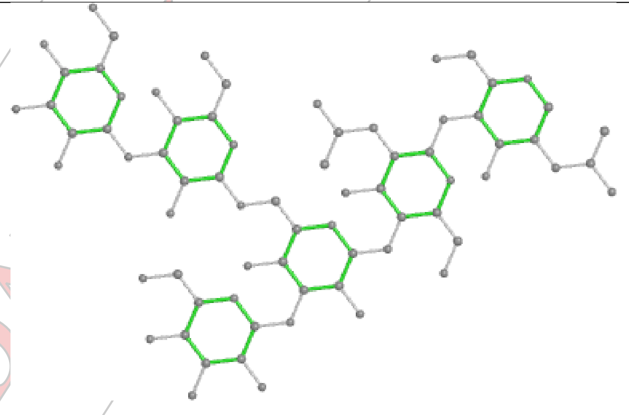

**Rings**

**Oligosaccharide Chain F**

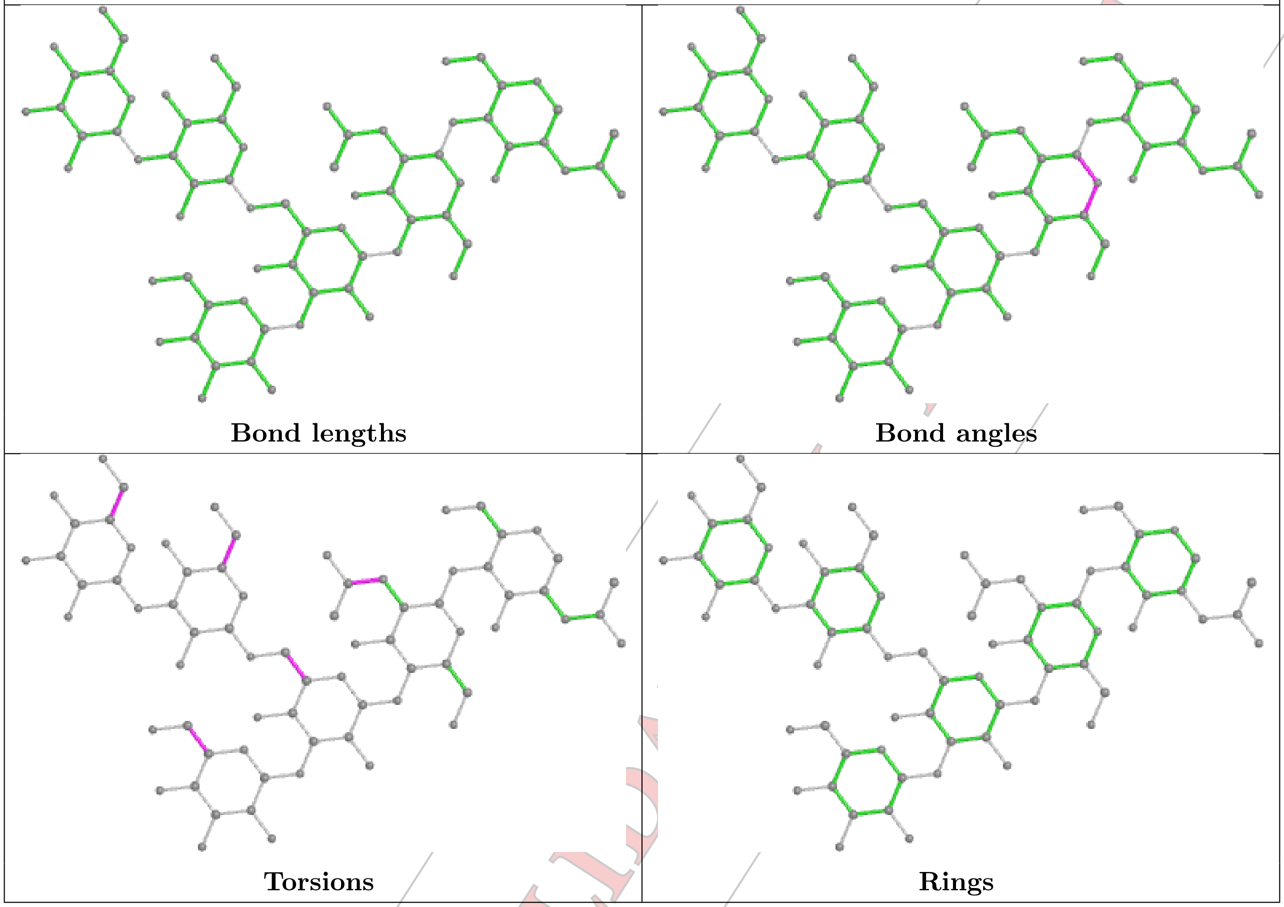


#### 5.6 Ligand geometry (i)

3 ligands are modelled in this entry.

In the following table, the Counts columns list the number of bonds (or angles) for which Mogul statistics could be retrieved, the number of bonds (or angles) that are observed in the model and the number of bonds (or angles) that are defined in the Chemical Component Dictionary. The Link column lists molecule types, if any, to which the group is linked. The Z score for a bond length (or angle) is the number of standard deviations the observed value is removed from the expected value. A bond length (or angle) with $|Z| > 2$ is considered an outlier worth inspection. RMSZ is the root-mean-square of all Z scores of the bond lengths (or angles).

| Mol | Type | Chain | Res | Link | Bond lengths | | | Bond angles | | |
|---|---|---|---|---|---|---|---|---|---|---|
| | | | | | Counts | RMSZ | $\#\|Z\| > 2$ | Counts | RMSZ | $\#\|Z\| > 2$ |
| 3 | NAG | B | 900 | 1 | 14,14,15 | 0.25 | 0 | 17,19,21 | 0.44 | 0 |
| 3 | NAG | A | 900 | 1 | 14,14,15 | 0.27 | 0 | 17,19,21 | 0.45 | 0 |
| 3 | NAG | C | 900 | 1 | 14,14,15 | 0.26 | 0 | 17,19,21 | 0.46 | 0 |

In the following table, the Chirals column lists the number of chiral outliers, the number of chiral centers analysed, the number of these observed in the model and the number defined in the Chemical Component Dictionary. Similar counts are reported in the Torsion and Rings columns.

'-' means no outliers of that kind were identified.

| Mol | Type | Chain | Res | Link | Chirals | Torsions | Rings |
|---|---|---|---|---|---|---|---|
| 3 | NAG | B | 900 | 1 | - | 2/6/23/26 | 0/1/1/1 |
| 3 | NAG | A | 900 | 1 | - | 2/6/23/26 | 0/1/1/1 |
| 3 | NAG | C | 900 | 1 | - | 2/6/23/26 | 0/1/1/1 |

There are no bond length outliers.

There are no bond angle outliers.

There are no chirality outliers.

All (6) torsion outliers are listed below:

| Mol | Chain | Res | Type | Atoms |
|---|---|---|---|---|
| 3 | B | 900 | NAG | C4-C5-C6-O6 |
| 3 | A | 900 | NAG | C4-C5-C6-O6 |
| 3 | C | 900 | NAG | C4-C5-C6-O6 |
| 3 | B | 900 | NAG | O5-C5-C6-O6 |
| 3 | A | 900 | NAG | O5-C5-C6-O6 |
| 3 | C | 900 | NAG | O5-C5-C6-O6 |

There are no ring outliers.

No monomer is involved in short contacts.

#### 5.7 Other polymers (i)

There are no such residues in this entry.

#### 5.8 Polymer linkage issues (i)

There are no chain breaks in this entry.

PRELIMINARY VALIDATION REPORT

### 6 Map visualisation (i)

This section contains visualisations of the EMDB entry D_1000251554. These are intended to permit visual inspection of the internal detail of the map and identification of artifacts.

#### 6.1 Orthogonal projections (i)

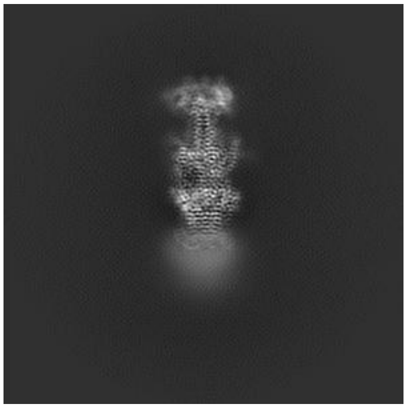
X

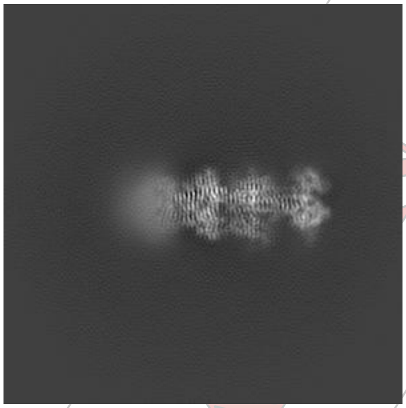
Y

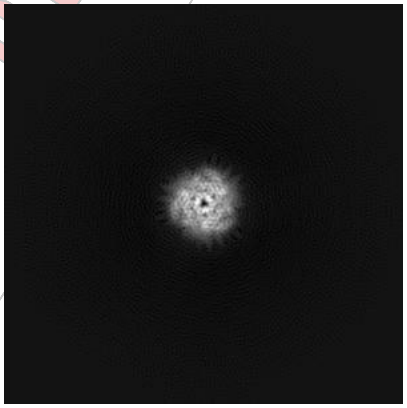
Z

The images above show the map projected in three orthogonal projections, in greyscale.

#### 6.2 Central slices (i)

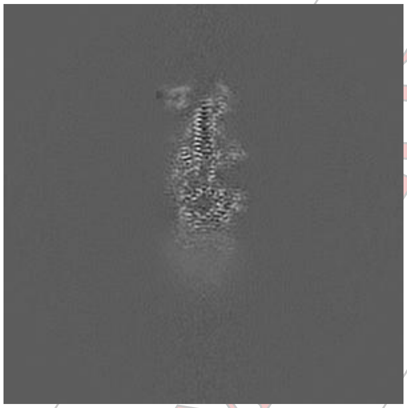
X

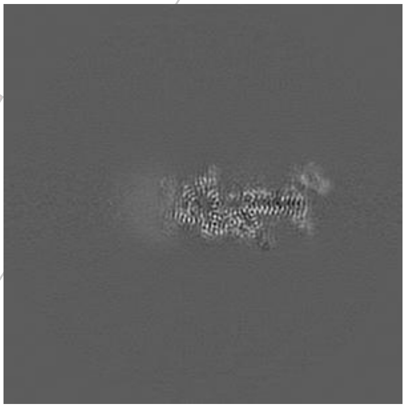
Y

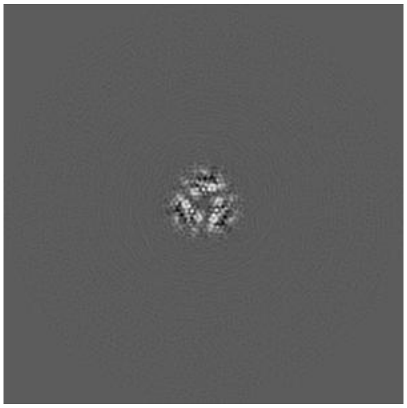
Z

The images above show central slices of the map in three orthogonal directions, in greyscale.

PRELIMINARY VALIDATION REPORT

#### 6.3 Largest variance slices (i)

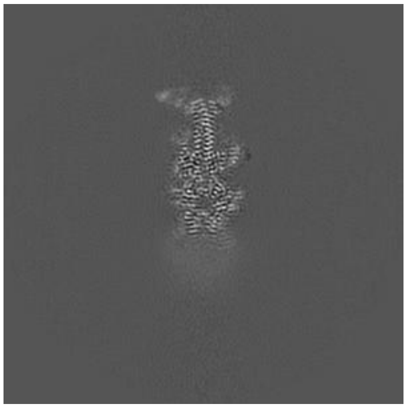

X Index: 205

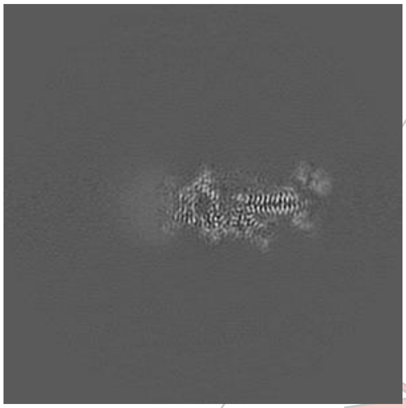

Y Index: 203

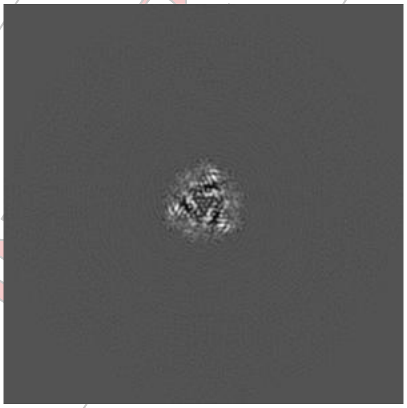

Z Index: 207

The images above show the highest variance slices of the map in three orthogonal directions, in greyscale.

#### 6.4 Orthogonal surface views (i)

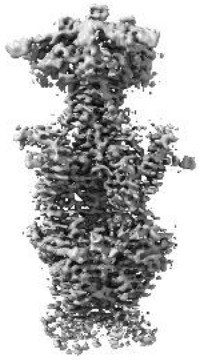

X

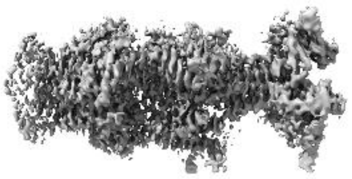

Y

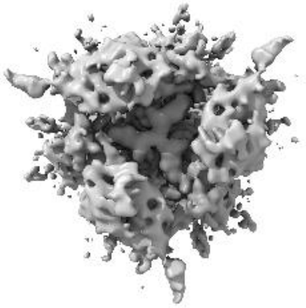

Z

The images above show the 3D surface view of the map at the recommended contour level 0.02. This in conjunction with the slice images can indicate whether an appropriate contour level has been selected.

#### 6.5 Mask visualisation (i)

This section was not generated. No masks were provided.

PRELIMINARY VALIDATION REPORT

### 7 Map analysis (i)

This section contains the results of statistical analysis of the map.

#### 7.1 Map-value distribution (i)

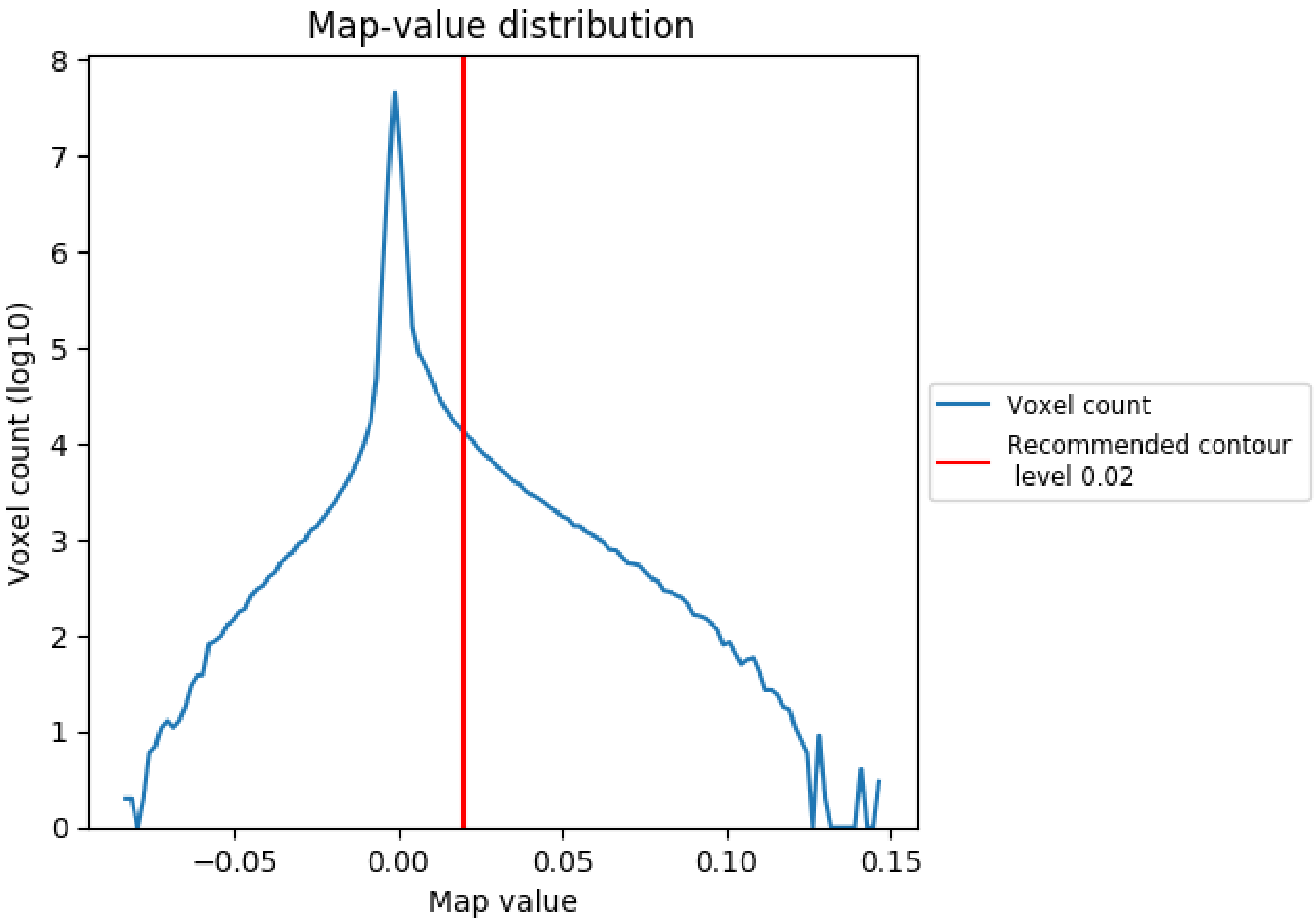


The map-value distribution is plotted in 128 intervals along the x-axis. The y-axis is logarithmic. A spike in this graph at zero usually indicates that the volume has been masked.

PRELIMINARY REPORT

#### 7.2 Volume estimate (i)

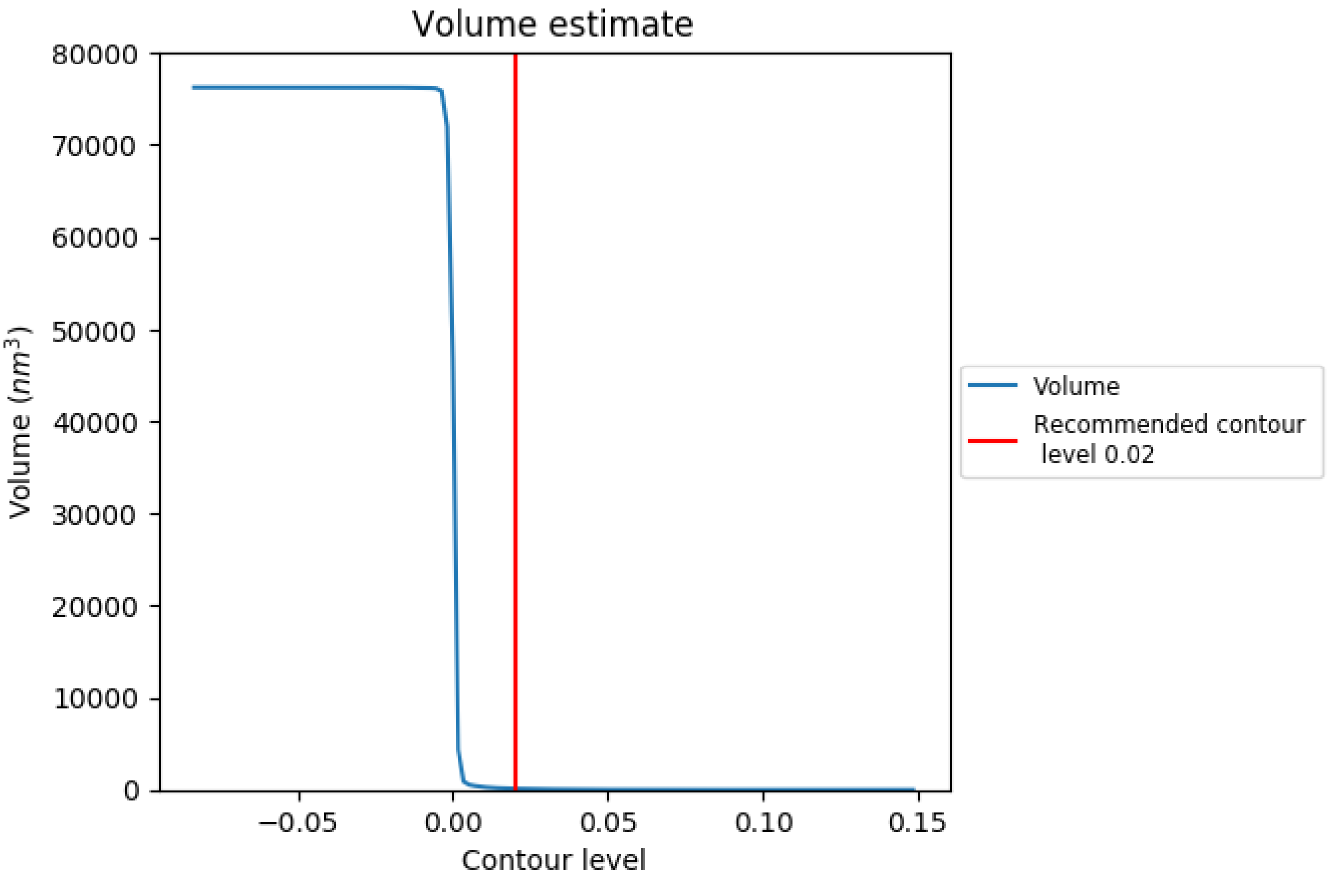


The volume at the recommended contour level is 129 nm$^3$; this corresponds to an approximate mass of 117 kDa.

The volume estimate graph shows how the enclosed volume varies with the contour level. The recommended contour level is shown as a vertical line and the intersection between the line and the curve gives the volume of the enclosed surface at the given level.

PRELIMINARY VA

#### 7.3 Rotationally averaged power spectrum (i)

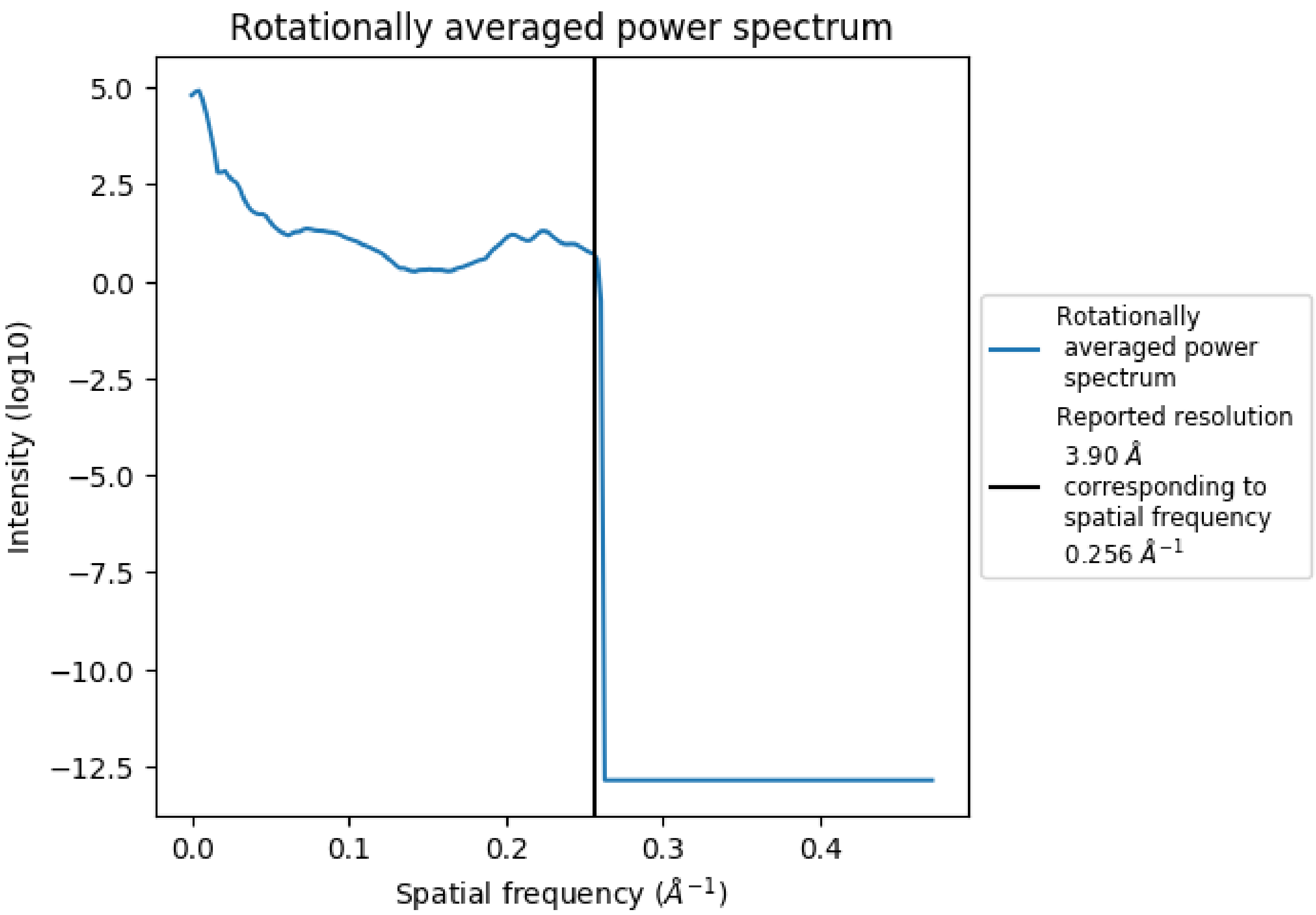


PRELIMINARY VALIDATION REPORT

### 8 Fourier-Shell correlation (i)

Fourier-Shell Correlation (FSC) is the most commonly used method to estimate the resolution for single-particle and subtomogram-averaging methods. The shape of the curve depends on the imposed symmetry, mask and whether or not the two 3D reconstructions used were processed from a common reference. The reported resolution is shown as a black line. Curves are displayed for $3\sigma$, 1-bit and 1/2-bit in addition to lines showing the 0.143 gold standard cut-off, 0.333 cut-off and legacy 0.5 cut-off.

#### 8.1 Resolution estimates (i)

These are global values for the map.

| Source | Criterion | Resolution estimate (Å) |
|---|---|---|
| Reported value | Not provided | 3.90 |
| Author-provided FSC | FSC 0.5 CUT-OFF | 4.11 |
| Author-provided FSC | FSC 1 BIT CUT-OFF | 3.96 |
| Author-provided FSC | FSC 0.33 CUT-OFF | 3.96 |
| Author-provided FSC | FSC 1/2 BIT CUT-OFF | 3.87 |
| Author-provided FSC | FSC 0.143 CUT-OFF | 3.86 |
| Author-provided FSC | FSC 3 SIGMA CUT-OFF | 3.79 |

#### 8.2 Calculated FSC (i)

This section was not generated. Half-maps were not provided.

PRELIMINARY VALIDATION REPORT

#### 8.3 Author-provided FSC (i)

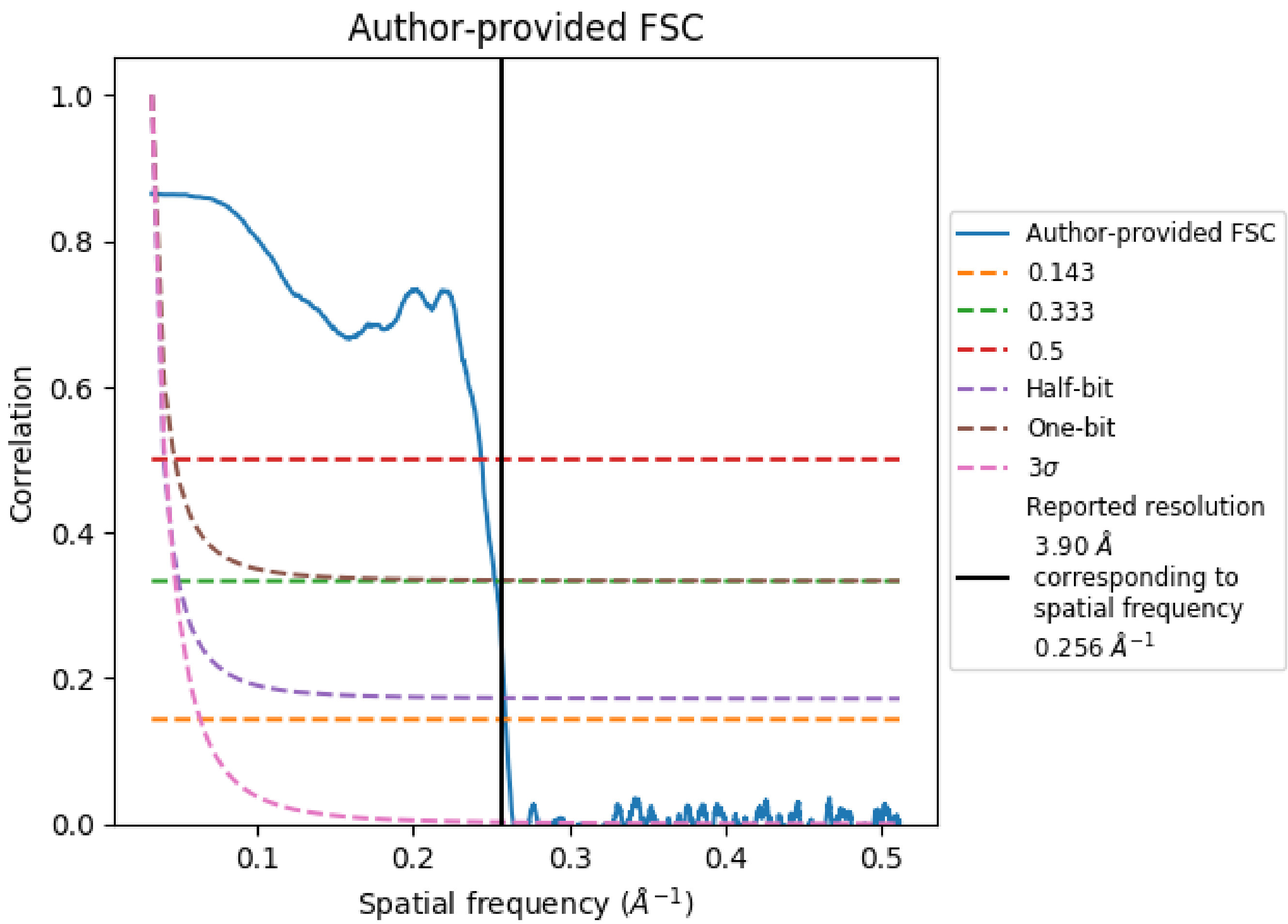


This FSC information was provided by the depositor.

PRELIMINARY

### 9 Map-model fit (i)

This section contains information regarding the fit between EMDB map D_1000251554 and PDB model D_1000251554. Per-residue inclusion information can be found in section 3 on page 5.

#### 9.1 Map-model overlay (i)

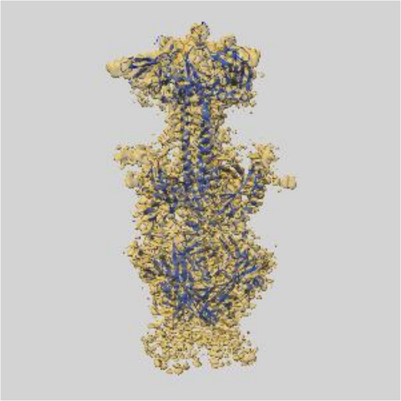

X

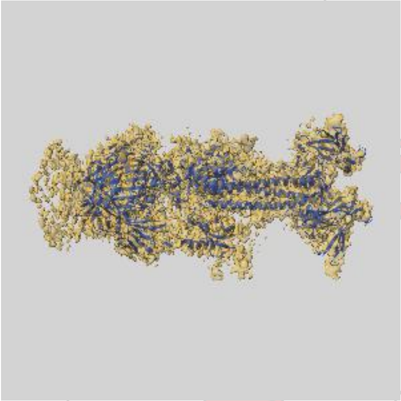

Y

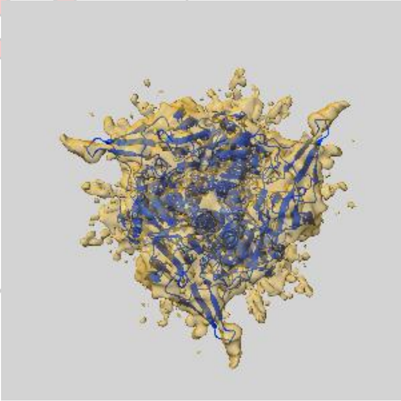

Z

The images above show the 3D surface view of the map at the recommended contour level 0.02 at 50% transparency in yellow overlaid with a ribbon representation of the model coloured in blue. These images allow for the visual assessment of the quality of fit between the atomic model and the map.

PRELIMINARY VALIDATION REPORT

#### 9.2 Atom inclusion (i)

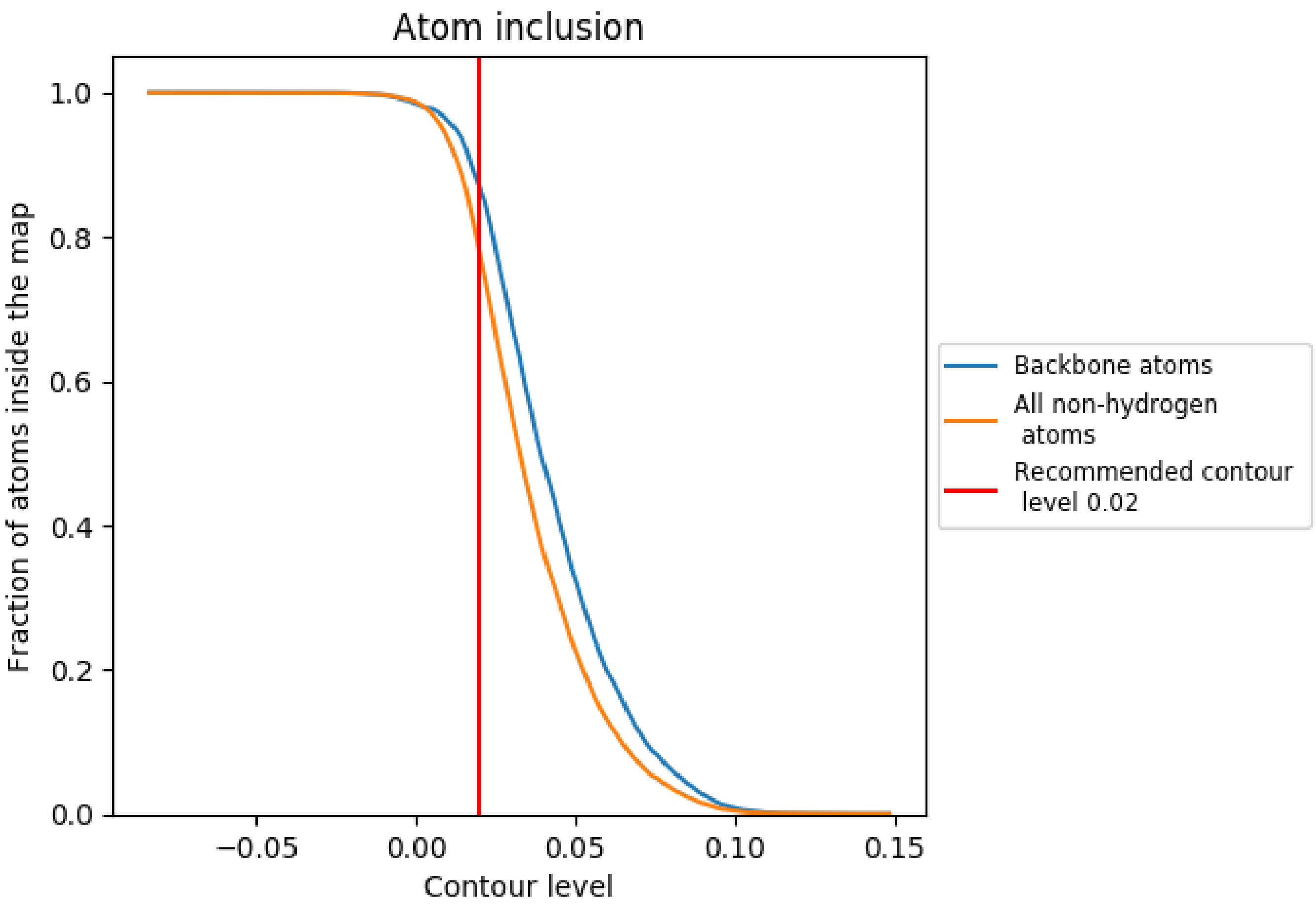


At the recommended contour level, 87% of all backbone atoms, 78% of all non-hydrogen atoms, are inside the map.

PRELIMINARY
